# Repurposing of the RIPK1 selective benzo[1,4]oxazepin-4-one scaffold for the development of a type-III LIMK1/2 inhibitor

**DOI:** 10.1101/2025.02.03.636296

**Authors:** Sebastian Mandel, Thomas Hanke, Sebastian Mathea, Deep Chatterjee, Hayuningbudi Saraswati, Benedict-Tilman Berger, Martin-Peter Schwalm, Satoshi Yamamoto, Michiko Tawada, Terufumi Takagi, Sandra Röhm, Ana Corrionero, Patricia Alfonso, Maria Baena, Lewis Elson, Amelie Menge, Andreas Krämer, Raquel Pereira, Susanne Müller, Daniela S. Krause, Stefan Knapp

## Abstract

Benzoxazepinones have been extensively studied as exclusively selective RIP kinase 1 inhibitors. This scaffold binds as a type-III inhibitor targeting the αC-out/DFG-out conformation. This inactive conformation results in a large expansion of the kinase back pocket, a conformation that has also been reported for LIM kinases. Scaffold hopping is common in the design of orthosteric kinase inhibitors, but has not been explored in the design of allosteric inhibitors, mainly due to the typically exclusive selectivity of type III inhibitors. Here, we hypothesized that the shared structural properties of LIMKs and RIPKs could lead to novel type III LIMK inhibitors using the benzoxazepinone scaffold. We report the discovery of a novel LIMK1/2 inhibitor that relies on this scaffold-based approach. The discovered compound **10** showed low nanomolar potency on LIMK1/2 and exceptional selectivity, as confirmed by a comprehensive selectivity panel with residual RIPK activity as the only off-target. The study provides one of the few examples for scaffold hopping for type-III inhibitors which are usually associated with exclusive target selectivity.

**Table of contents (TOC) graphics:** 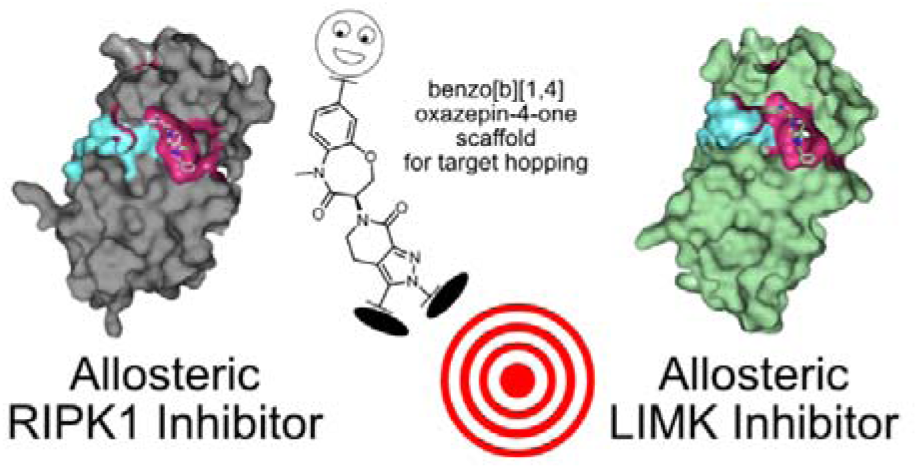

## Introduction

Protein kinases form a large family of proteins with largely conserved ATP binding sites. Their pivotal role in the development of diseases led to their extensive exploitation as drug targets. To date, protein kinases are one of the most successful protein families for the development of new medicines.^1–3^ However, the selectivity challenge in the development of ATP-competitive inhibitors has limited the discovery of new kinase inhibitors to the field of oncology for many years. The reason for this is that the promiscuity of inhibitors can be tolerated during short treatment cycles or, due to the activation of multiple signaling pathways in cancer, the broad-spectrum activity of inhibitors can even be an advantage for therapy. Several strategies have evolved that may lead to selective kinase inhibitors, comprising shape complementarity of ATP mimetic compounds^4,5^, covalent kinase inhibitors^6,7^ and targeting unique conformations of the catalytic domain ^8,9^.

One of the most efficient strategies achieving exclusive selectivity is often only possible by developing allosteric inhibitors that bind to the kinase back pocket formed by an outward shift of the αC helix (type-III) or to other allosteric pockets that may be located anywhere on the surface of the catalytic kinase domain (type-IV). Excellent examples of such inhibitors are type-III MEK1/2 inhibitors,^10,11^ as well as allosteric inhibitors of ABL1 and ABL2^12^ that bind to the myristate binding pocket or compounds that address the αD pocket of CK2 and thus exhibit exclusive selectivity.^13^ However, the exclusive selectivity of allosteric kinase inhibitors hampers broader development of allosteric inhibitors, as scaffold hopping, which is extensively used for ATP mimetic compounds, cannot be readily applied to allosteric inhibitors.^14^

The receptor-interacting serine/threonine-protein kinase 1 (RIPK1) is a key regulator mediating the delicate balance between pro-survival signaling and cell death in response to a broad set of inflammatory stimuli. Its broad implication in inflammatory diseases, neurodegenerative processes, ischemia and acute inflammatory conditions, such as sepsis positioned RIPK1 into the focus of drug research.^15^ Early drug development efforts identified Necrostatins (**1**), a widely used potent and selective RIPK1 inhibitor, that targets an allosteric pocket induced by the αC-out and DFG-out inactive states of RIPK1.^16^ Later, a DNA-encoded library screen and chemical optimization identified GSK’481 (**2**)^17^, a highly potent and mono-selective RIPK1 inhibitor harboring a benzoxazepinone scaffold that was further developed into the clinical candidate GSK2982772 (**3**), which has been investigated in phase 2 clinical trials for psoriasis, rheumatoid arthritis and ulcerative colitis.^18^ The benzoxazepinone scaffold has been extensively exploited for the development of allosteric RIPK inhibitors resulting for instance in GNE684 (**4**) by Genentech^19^, Eclitasertib (DNL-758) (**5**) developed by Delali (WO2017136727A2) and the brain penetrant compound **22** a derivative of the RIPK1 probe (TP-030-1), by Takeda (**6**) (**Fig. 1**).^20^

**Figure 1:**
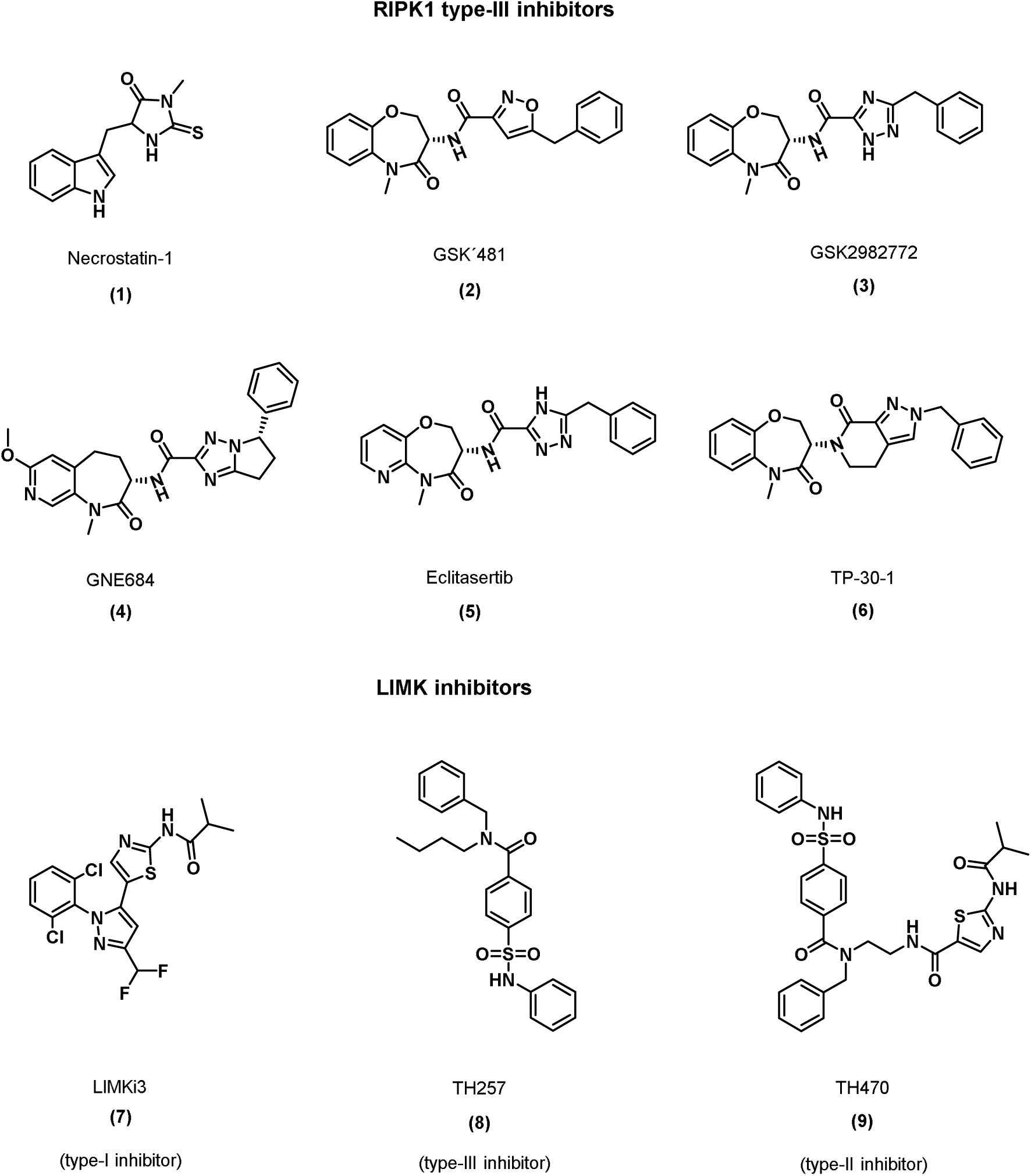
Allosteric inhibitors of RIPK1 and LIMK1/2 and commonly used tool compounds.

While all these inhibitors are exclusively selective, we hypothesized that the benzoxazepinone scaffold could also be adapted to allosterically inhibit protein kinases that are known to assume the αC-out and DFG-out inactive states and focused our attention on the related TKL (tyrosine kinase like) family members LIMK1/2 (LIM Domain Kinase 1/2). LIMKs are primarily known as effectors of the small GTPases of the Rho family pathway that activated Rho kinases (ROCK) or p21-activated kinases (PAKs) regulating cell motility. Deregulation of LIMKs has been reported in various diseases such as in cancer or glaucoma, which is why a number of inhibitors have been developed.^21–23^ Among the small molecules developed are also allosteric inhibitors that bind to the αC-out and DFG-out pocket (**7**, **8**, **9**).^24,25^ The type-III inhibitor TH257, and the chimeric type-II inhibitor TH470, designed by adduct formation of the type-I inhibitor LIMKi3 and TH257 are highly potent and selective chemical probes that have been demonstrated to be efficacious in cellular models of LIMK1 associated diseases such as Fragile X syndrome caused by loss-of-function mutation of the fragile X mental retardation 1 (*FMR1*) gene.^26^ Comparison of the inactive states of RIPK1 and LIMK αC-out and DFG-out conformation suggests that the benzoxazepinone scaffold could be adapted inhibiting also LIMK1/2. Screening of type-III inhibitors targeting RIPK1 identified LIJTF500025 (**10**), a potent allosteric LIMK1/2 and RIPK1 dual inhibitor that represents a template for benzoxazepinone-based LIMK inhibitors with excellent pharmacological properties. Here we describe the characterization of **10** as a chemical tool compound for LIM kinases.

## Results and Discussion

Our hypothesis that the scaffold of the benzoxazepinone-based type-III RIPK1 inhibitor targets LIM kinases, was based on a similar conformation of the inactive type-III binding state as observed by TH257 and TH300.^25^ The binding modes of these compounds are based on a shared αC-out and DFG-out conformation, which has also been observed in crystal structures of benzoxazepinone-based RIPK1 inhibitors.^25,27^ Representative members of both TKL subfamilies such as LIMK2 and RIPK1, exhibit a sequence identity of 72.6% within their catalytic domains. Comparison of co-crystal structures (**Fig. 2**) of both proteins in complex with the corresponding type-III inhibitors confirmed the conservation of binding of allosteric type-III inhibitors to both target families suggesting that benzoxazepinone-based inhibitors may also inhibit LIM kinases.

**Figure 2:**
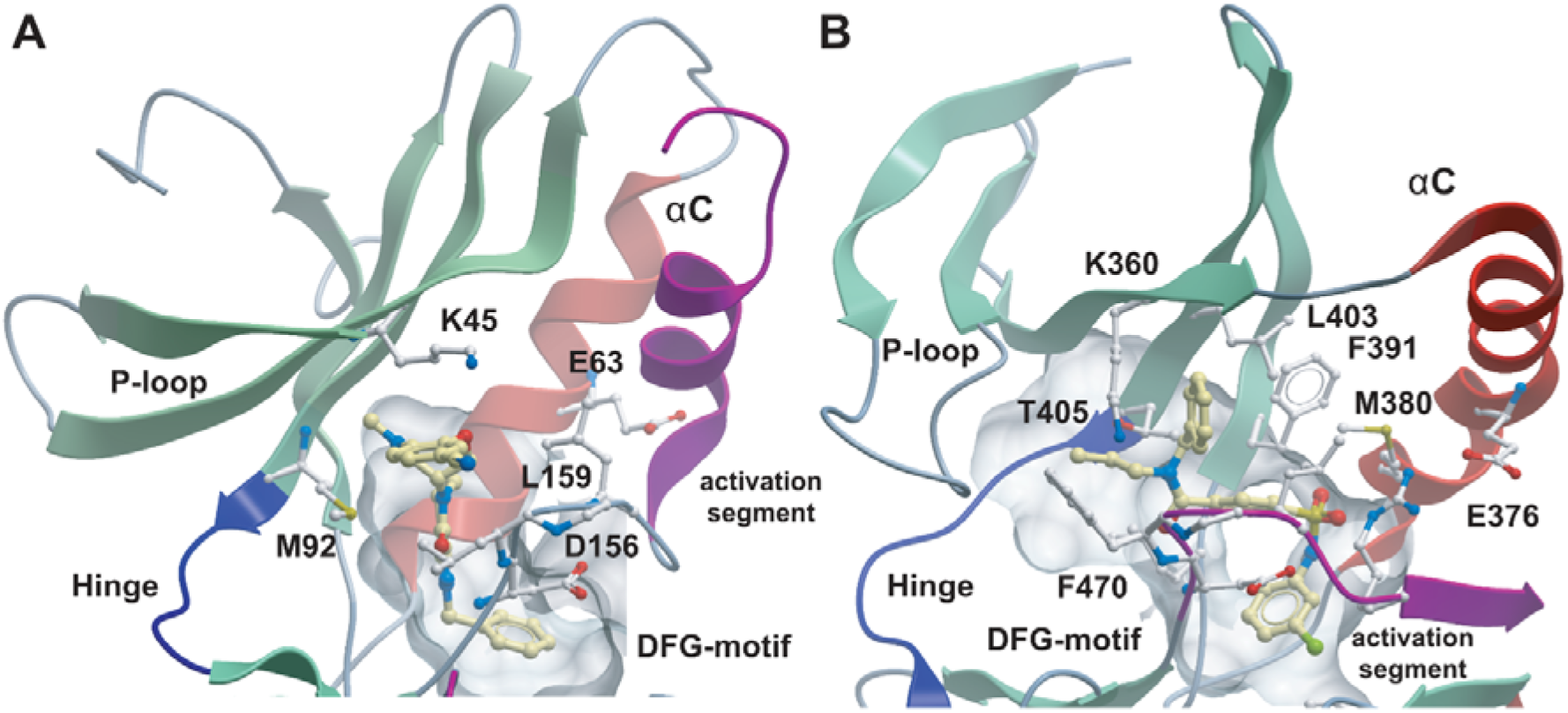
Comparison of RIPK1 (**A**) and LIMK2 (**B**) type-III inhibitor complexes (RIPK1-GSK481 (5HX6) and LIMK2-TH300 (5NXD). Inhibitors are highlighted by yellow carbon atoms. The backbone of the kinase hinge region is highlighted in blue, αC in red and the DFG-motif as well as the activation segment is colored in purple. Main interacting residues are shown in ball and stick representation and are labelled.

In the structure shown, GSK481 binds deeply into the back pocket and does not interact with the hinge backbone (**Fig. 2**). The amide carbonyl moiety attached to the isoxazole forms a direct hydrogen-bond with the backbone amide nitrogen of Asp156. Compared to the type-I binding mode, the αC-helix exhibited an outwardly oriented conformation, while the activation loop in the RIPK1/GSK481 complex was structured.^17^ Consistent with this binding mode, no salt bridge was present between Lys45 and Glu63, which is a hall mark of the inactive kinase state. Similar features were observed in the LIMK2/TH300 complex, in which the αC-helix was distant from the ATP binding site, with a distance of 13,8 Å between the VIAK-motif K360 and αC E376, the two residues that form a salt bridge in active kinases (**Fig. 2**). ^28^ Furthermore, a significant distortion of the phosphate binding loop (P-loop) was observed with an inward flip of F470, suggesting that these three structural elements (DFG, P-loop and αC) exhibit substantial flexibility in the unphosphorylated state of LIMK2.

TH257 is a highly selective LIMK1/2 inhibitor but compounds of this series have been associated with poor metabolic stability, limiting the utility of TH257 to *in vitro* studies. In contrast, benzoxazepinones have excellent metabolic stability, which has led to inhibitors being used in clinical studies.^18^ To test whether allosteric RIPK inhibitors bind and inhibit LIM kinases, we screened a small library of benzoxazepinones that have been developed by Takeda. All compounds contained the 7-oxo-2,4,5,7-tetrahydro-6H-pyrazolo[3,4-c]pyridine core as reported by Yoshikawa et al.^20^ Since these inhibitors interact with the inactive state, we used differential scanning fluorimetry (DSF) binding assays^29^ to identify hit compounds.

This effort identified compound **10** (LIJTF500025), ((*S*)-2-benzyl-6-(8-chloro-5-methyl-4-oxo-2,3,4,5-tetrahydrobenzo[*b*][1,4] oxazepin-3-yl)-7-oxo-4,5,6,7-tetrahydro-2*H*-pyrazolo[3,4-c]pyridine-3-carboxamide), as a ligand that strongly stabilized LIMK1 by 7.0 K and LIMK2 by 16.3 K in the DSF assay. In the same screen, we also identified the regio isomer LIJTF500120a (**11**), which was inactive against LIMK1 and LIMK2, suggesting that this closely related analog may serve as a negative control compound. Compared to the highly selective RIPK1 inhibitor TP-30-1 (**6**), the 5-methyl-2,3-dihydro-1,5 benzooxazepine-4-one core was surprisingly similar having only an additional chlorine at position 8 of the benzoazepine as well as the carboxamide attached to the pyrazole moiety. A closely related RIPK1 inhibitor (**12**) containing a nitrile at position 8 and a chlorine instead of the carboxamide moiety in **10**, has been co-crystallized with the catalytic domain of RIPK1 (pdb-ID: 6C4D).^20^ The binding mode of this type-III inhibitor in RIPK1 revealed that the nitrile points towards the solvent and does not interact with the RIPK1 catalytic domain. However, the chlorine moiety on the pyrazole ring was directed towards the αC residue M67 and large moieties at this position of the inhibitor would therefore be expected to lower affinity for RIPK1.

In order to confirm the binding mode of **10** in LIM kinases, we crystallized compound **10** with LIMK1 (pdb-ID: 7ATU). The structure was refined at 2.8 Å with clear electron density for the ligand. As expected, compound **10** bound to the back pocket of the catalytic domain created by the αC-out and DFG-out conformation as described for previous type-III LIMK inhibitors (**Fig. 3B**).^25^ The inhibitor was stabilized by a number of mainly aromatic interactions with the upper lobe hydrophobic core structure. The structure explained the inactivity of the regioisomer **11**, which would clash with the lower lobe of the kinase catalytic domain. Comparison with the RIPK complex (pdb-ID6c4d) revealed amazing conservation of the binding mode of both inhibitors (**Fig 3C, 3D**).

**Figure 3:**
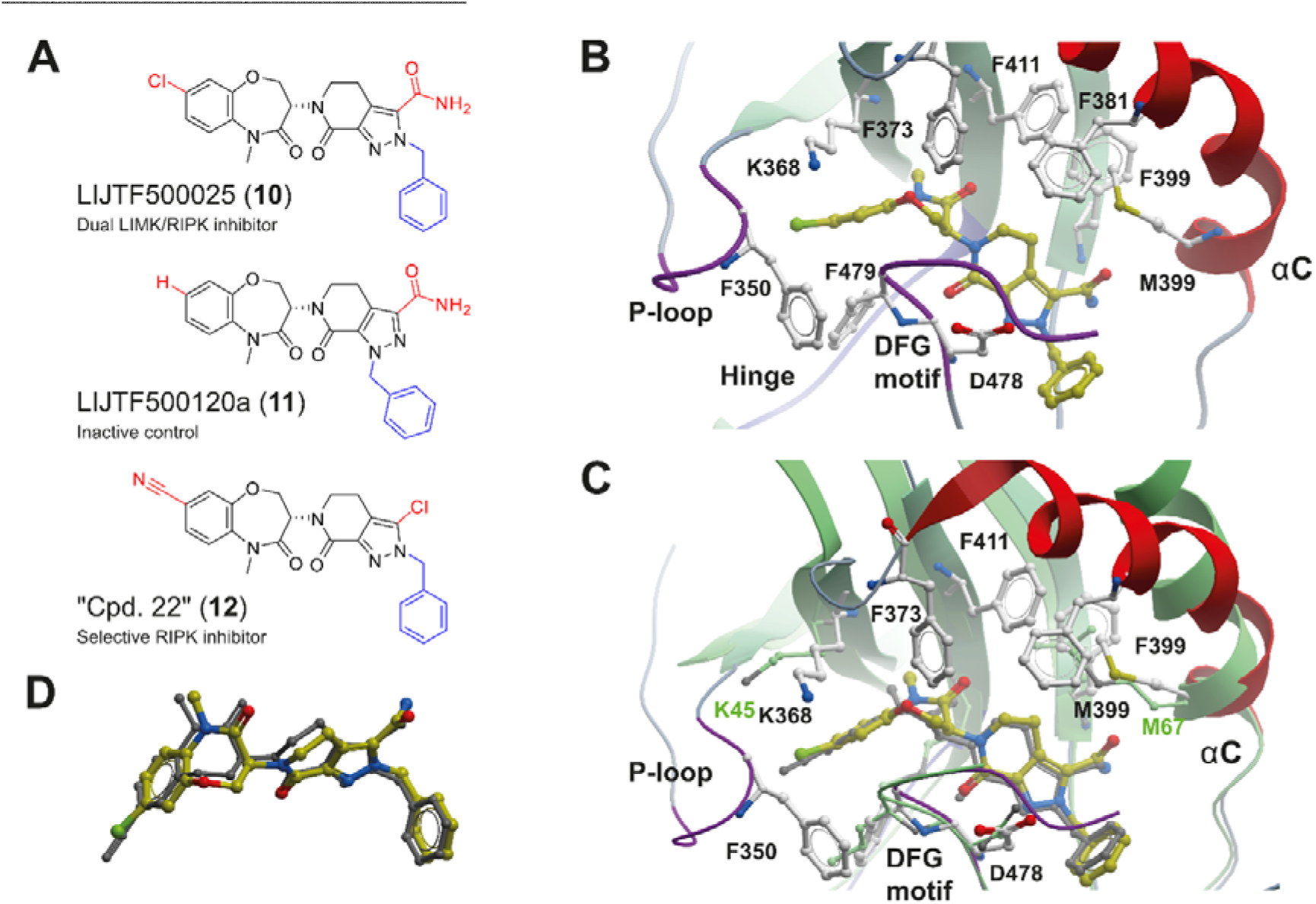
**A**) Chemical structure of the lead benzoxazepinone (**10**) and the corresponding inactive derivative (**11**). **B**) Structure of compound **10** in complex with LIMK1 (pdb-ID: 7ATU). Compound **10** is highlighted with yellow carbon atoms in ball and stick representation. The main interacting residues are shown and structural elements have been labelled. **C**) Superimposition of the LIMK1 compound **10** structure with compound 22 (**12**) from Yoshikawa et al. (pdb-ID: 6c4d) in RIPK1. **D**) Detailed view of the superimposition of both inhibitors **10** and **12** extracted from the corresponding crystal structures.

Next we resynthesized compound **10** according to published procedures. LIJTF500025 (**10**) was synthesized in an 8-step synthesis (**Scheme 1**). The synthesis of the building blocks **18** and **19** was performed as previously described in the literature^17,30^ and the patent (WO2006061136A3). In brief, we first synthesized building block **18** according to scheme 1. The synthesis was carried out starting from commercial diethyl but-2-ynedioate (**13**), which was reacted with benzylhydrazine (**14**) to obtain the corresponding pyrazole (**15**). Pyrazole **16** was obtained by halogenation and Vilsmeier-Haack reaction with phosphorus oxybromide and DMF. Subsequent Wittig reaction with methoxymethyl triphenylphosphonium chloride provided compound **17** and ether cleavage under acidic conditions with tautomerization led to building block **18**.

**Scheme 1:**
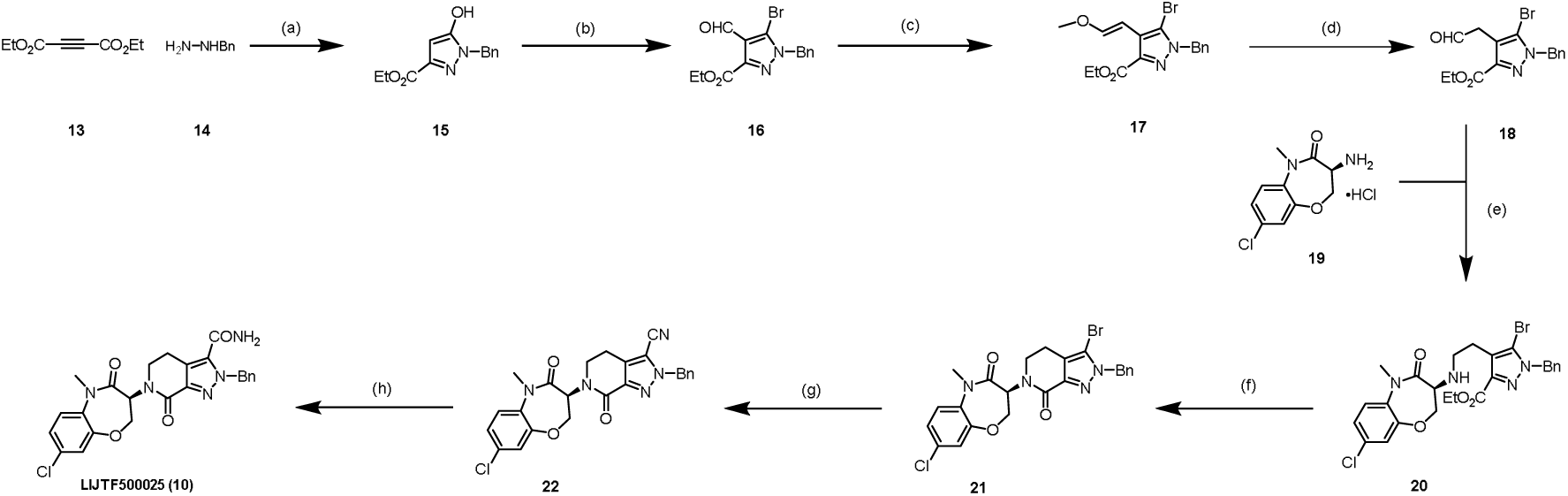
Synthetic route of compound 10. Reagents and conditions: (a) K_2_CO_3_, EtOH, 16 h, 90 °C; (b) POBr_3_, DMF, DCE, 16 h, 90 °C; (c) Ph_3_P^+^CH_2_OMeCl^-^, KOtBu, THF, 16 h, 0–15 °C; (d) aq. HCl, THF, 16 h, 15∼60°C, (e) α-picoline-borane, AcOH, MeOH, 2 h, 0–20 °C; (f) AlMe_3_, toluene, 1 h, 0∼100 °C; (g) Zn(CN)_2_, Pd(PPh_3_)_4_, DMF, 4 h, 100 °C; (h) H_2_O_2_, K_2_CO_3_, DMSO, 1 h, 20 °C.

Building block **18** was then coupled via a reductive amination with building block **19** to the secondary amine (**20**). Lactamization led to compound **21**, which was then subjected to palladium-catalyzed cyanation to obtain **22**. Hydrolysis of the nitrile **22** under basic conditions using hydrogen peroxide yields to LIJTF500025 (**10**).

The negative control compound **11** was synthesized according to **scheme 2**. Reductive amination of **23** and **18** yielded the secondary amine **24**. Lactamization provided compound **25** and palladium-catalyzed cyanation let to nitrile **26**, which was hydrolyzed to primary amide **27**. Palladium-catalyzed hydrogenation resulted in the debenzylated compound **28** that was then benzylated with benzyl chloride to the neg. control compound **LIJTF500120** (**11**)

**Scheme 2:**
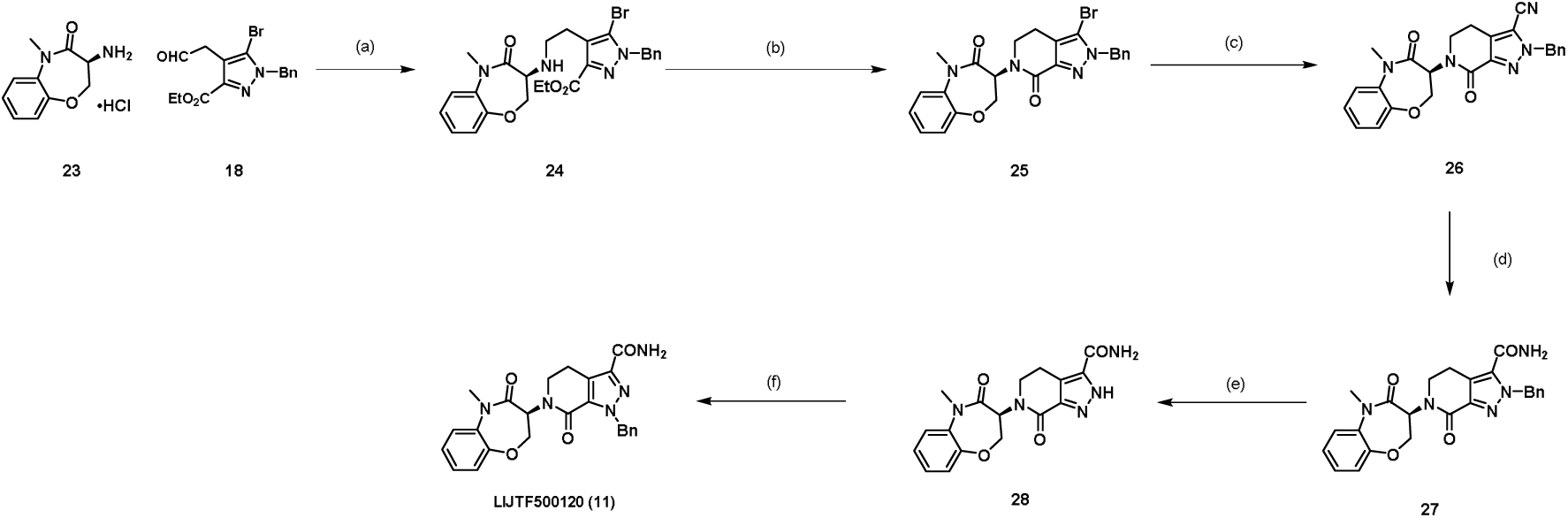
**Synthetic route of compound 11**: Reagents and conditions: (a) boran-2-methylpyridin, AcOH, MeOH, 2 h, 0 - 20 °C; (b) AlMe_3_, toluene, DCE, 1 h, 0 - 100 °C; (c) Zn(CN)2, Pd(PPh_3_)_4_, DMF, 4 h, 100 °C; (d) H_2_O_2_, K_2_CO_3_, DMSO, 1 h, 20 °C °C, (e) Pd(OH)_2_, H_2_, aq. HCl, MeOH, 12 h, 20 °C; (f) BnCl, Cs_2_CO_3_, DMF, 12 h, 80 °C

Primarily, we were interested in the selectivity of **10** and the inactive control compound **11**. Initially, we tested both compounds with an in-house panel of over 107 proteins in a DSF assay that included recombinant protein kinases and off-targets that are frequently inhibited by kinase inhibitors (**Fig 4A**). Analysis of the Δ*T*_m_ shifts revealed LIMK1 and LIMK2 as the only targets that were significantly stabilized by **10** (16.3 K for LIMK2 and 7.0 K for LIMK1). The only potential off-target was casein kinase 2A (CK2A2) with a Δ*T*_m_ 2.3 K indicating binding in the micromolar K_D_ range.^31^ Compound **11** showed no significant shift except for CK2A2 (2.2 K). Based on the observed selectivity profile, we analyzed compound **10** using the ScanMAX KINOMEscan assay platform (Eurofins Scientific), which covers 468 kinases including some pathogenic mutants. Selectivity data are provided as supplemental data.

**Figure 4:**
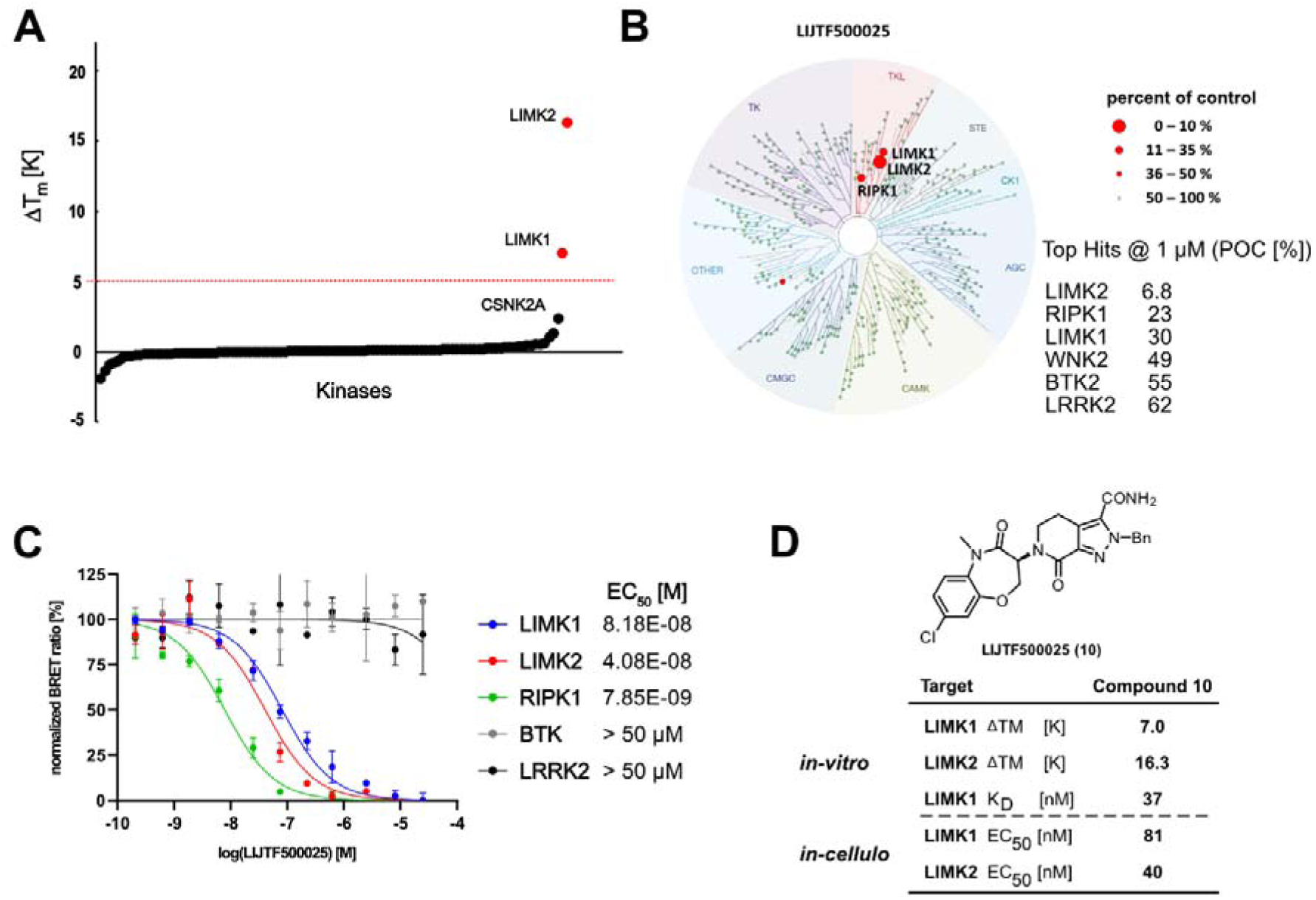
Selectivity of 10. **A**) Thermal shift selectivity profile of compound **10** towards a panel covering 107 proteins; **B**) Selectivity profile using the scanMAX kinome wide selectivity assay (Eurofins) of **10**. Data are illustrated using the TREEspot analysis; (**Table S6**) **C**) Target engagement and off-target evaluation based on the scanMAX data analyzed by NanoBRET assays. The EC_50_ values are provided in the graph legend. **D**) Assay data measured on the LIMK1/2 interaction with **10**. Shown are DSF data, ITC K_D_ data for LIMK1 as well as cellular target engagement data by NanoBRET.

Encouragingly, compound **10** demonstrated excellent selectivity for LIMK1/2 in this comprehensive panel and still activity for the original target RIPK1, with a selectivity score (S35) of 0.007 at a screening concentration of 1 μM. Weak activity was also observed for the kinases WNK2, BTK and LRRK2. Further selectivity was measured using a live-cell panel of 184 kinases based on the NanoBRET technology. In agreement with the thermal shift selectivity and the scanMAX kinome selectivity assay, only LIMK1, LIMK2 and RIPK were detected. For follow up of the detected hits, cellular on target activity was confirmed using NanoBRET^32^ revealing EC_50_ values of 81 nM and 40 nM for LIMK1 and LIMK2, respectively. Compound **10** had an EC_50_ value of 7.8 nM on RIPK1. No significant interaction was observed for the potential off-targets up to an inhibitor concentration of 50 μM (**Fig. 4C**). Thus, compound **10** was identified as a selective dual LIMK1/2 and RIPK1 inhibitor. In addition, we measured direct binding in solution on LIMK1 using ITC (**Fig. S1**) revealing a K_D_ value of 37 nM. Enzyme kinetic assays using rapid fire MS yielded in pIC_50_ values of 6.1 and 8.2, on LIMK1 and LIMK2, respectively.

Since for allosteric inhibitors it is known that they might have kinetic advantages over canonical type I inhibitors we evaluated the binding kinetic on LIMK1 of this novel type III inhibitor (**10**) and compared it with our previous allosteric chemical probe (TH257; **8**)^33^. Therefore, both compounds were evaluated in a TR-FRET binding kinetic assay (KINETICfinder) offered by Enzymlogic. These data revealed that LIJTF500025 (**10**) had a fast *k*_on_ (*k*_1_) and a slow *k*_off_ (*k*_2_) rate resulting overall in a prolonged on-target residence time for this novel allosteric inhibitor **10** (τ; see **Table 1** and **Fig. S2**). These findings were further supported by the affinity values obtained with KINETICfinder for both type III inhibitors, which show a strong correlation with the biochemical and cellular data presented here and in previous publications^25^.

**Table 1:** Kinetic profiling of both allosteric type III inhibitors: TH257 (8) and LIJTF500025 (10) on LIMK1 using KINETICfinder. *k*_1_: *k*_on_; *k*_2_: *k*_off_; τ: residence time; *K*_d_: dissociation constant.

| Compound | Target | $k_1$ (1/Ms) | $k_2$ (1/s) | $\tau$ (min) | $K_d$ (M) |
| --- | --- | --- | --- | --- | --- |
| TH257<br>(**8**)<br>(type-III) | LIMK1 | 2,44E+03 | 1,64E-03 | 1,02E+01 | 6,70E-07 |
| LIJTF500025<br>(**10**)<br>(type-III) | LIMK1 | 1,56E+04 | 7,48E-04 | 2,23E+01 | 4,80E-08 |

Next, we examined the activity of **10** on endogenous protein by Western blotting. For this experiment, we used the glioblastoma cell line LN-229. Western Blot analysis of the LIMK1/2 substrate cofilin showed weaker levels of phosphorylation already at a concentration of **10** of 100 nM. At 1 µM and higher concentrations, cofilin phosphorylation was almost completely abrogated. The negative control **11** showed no effect on cofilin phosphorylation at 5 µM. As a positive control compound we used the dual LIMK1/2 inhibitor LIMKi3^34^, which showed a similar activity on substrate phosphorylation (**Fig. 5**). These results indicate that the allosteric LIMK1/2 inhibitor **10** has a comparable potency in the cellular context as the type-I inhibitor.

**Figure 5:**
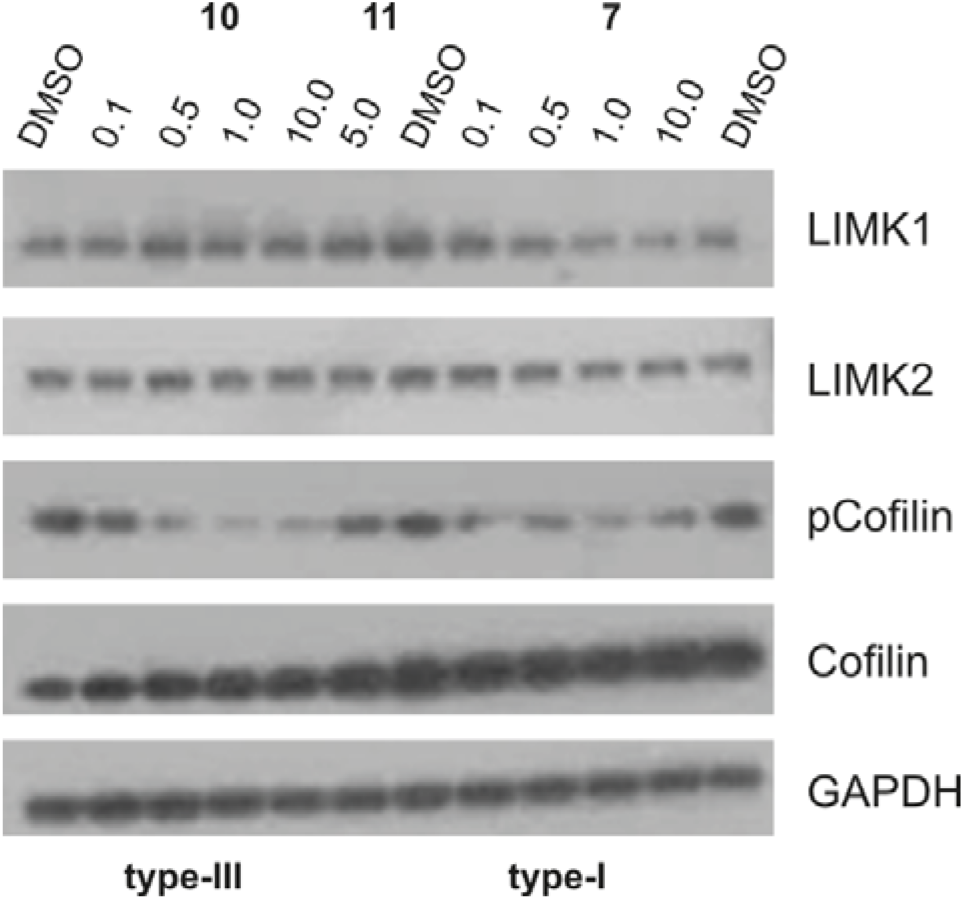
Western blot analysis. The LIMK inhibitors **10** and **7** (LIMKi3, a type-I inhibitor) and the negative control **11** were tested in Western blots using the LIMK substrate cofilin in LN-229 cell lysates. GAPDH was used as a loading control and total cofilin and LIMK1/2 levels were also assessed. Cells were treated for 6h with the inhibitors or the DMSO control.

Finally, toxicity of **10** and **11** were assessed using a multiplex toxicity and cellular health high content assay at concentrations of 1 μM and 10 μM, respectively. No general cytotoxicity was observed. The impact on tubulin, mitochondrial mass, and cellular permeability in healthy cells remained comparable to that of the 0.1% DMSO control (**Fig. 6**) for compound **10**. At a concentration of 1 μM, neither compound exhibited significant cytotoxicity across the tested cell lines. However, at the elevated concentration of 10 μM, a marginal increase in the apoptotic cell population was noted in HEK293T cells with prolonged incubation. Additionally, in U2OS cells, a minor, albeit non-significant, reduction in cell viability was observed upon treatment with 1 μM of the negative control, compound **11**.

**Figure 6:**
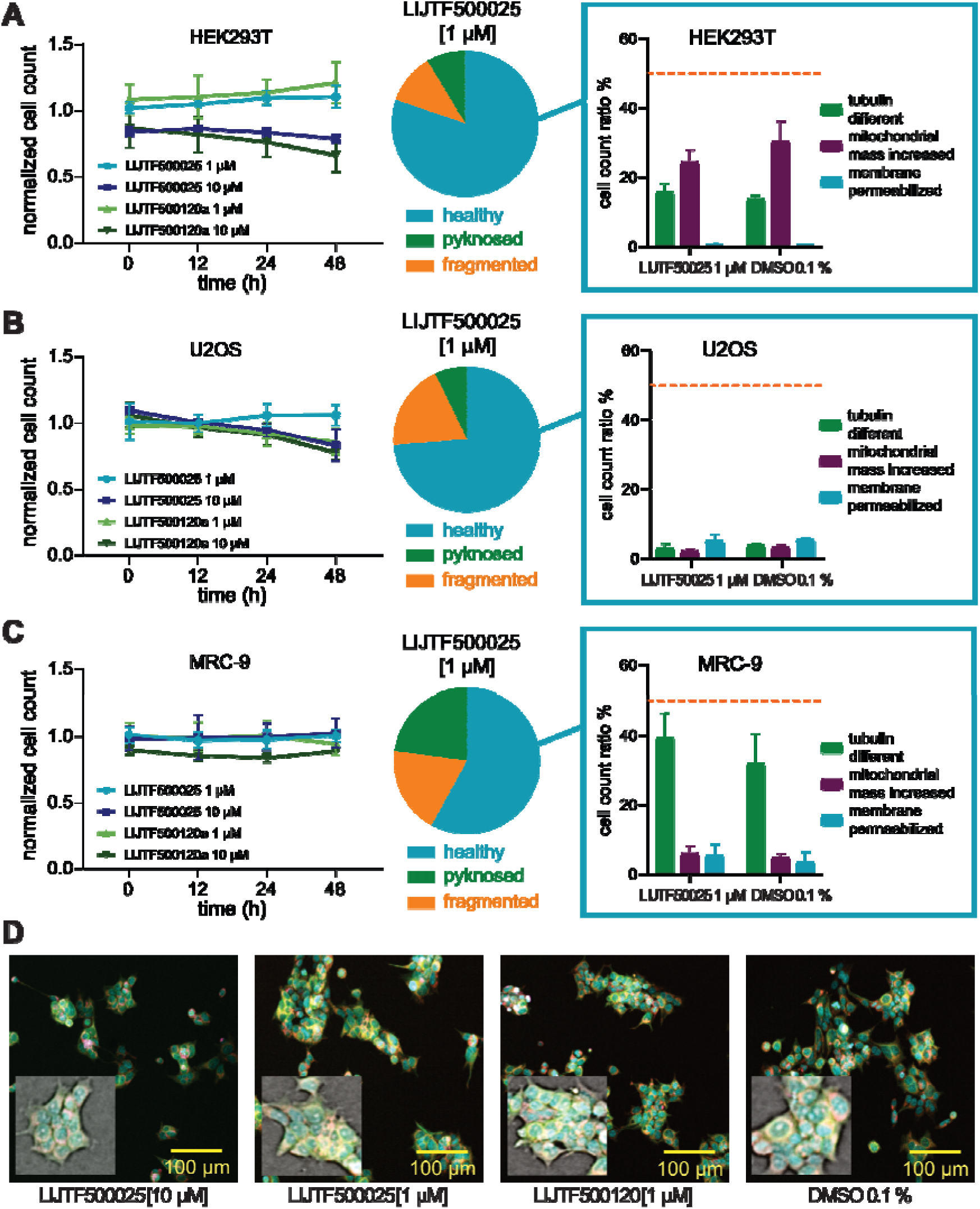
Live-cell viability assessment in HEK293T, U2OS and MRC-9 cells. Normalized cell count after 10 µM and 1 µM compound exposure (LIJTF500025, LIJTF500120a) measured over time (0 h, 12 h, 24 h, 48 h). Data were normalized to cells (HEK293T (**A**), U2OS (**B**) or MRC-9 (**C**)) exposed to DMSO (0,1 %). Fraction of healthy, pyknosed and fragmented nuclei of cells exposed to 1 µM of LIJTF500025 shown as pie charts. Fractions of cells showing a change in microtubule structure, having an increased mitochondrial mass or membrane permeabilization shown in comparison to cells exposed to 0.1 % DMSO as a negative control are highlighted. A threshold value of 50 % is marked in orange. Error bars show SEM of two biological replicates. **D** Fluorescent image and highlighted brightfield confocal image of stained (blue: DNA/nuclei, green: microtutubule, red: mitochondria, magenta: Annexin V apoptosis marker) HEK293T cells after a 24 h exposure to 10 µM and 1 µM compound (LIJTF500025, LIJTF500120a) in comparison to 0,1 % DMSO control. High content data are provided as supplemental data.

In summary, both LIJTF500025 (**10**) and LIJTF500120a (**11**) demonstrated no cytotoxicity at 1 μM across the evaluated cell lines. Only at the higher concentration of 10 μM there was a slight increase in toxicity detected within the 48-hour timeframe. However, due to the absence of LIJTF500025 being tested at 10 μM in the KINOMEscan, it remains inconclusive whether these observed effects are attributable to non-specific cytotoxicity or potential kinase inhibition at higher concentrations.

Encouraged by the cellular target engagement data by NanoBRET and the cellular activity on the phosphorylation levels of cofilin we were interested to study the pharmacokinetics of compound **10**. Bearing in mind that several benzoxazepinones went into clinical trials and with the knowledge that our previous allosteric type III LIMK1/2 inhibitor had a poor metabolic stability we first evaluated the metabolic stability of compound **10** in human/mouse/rat. As expected, the hepatic metabolic stability was about a magnitude better than our previous type III inhibitor **TH257** (Cl [µL/min/mg]; h/m/r 22/69/5 vs 470/>768/>768) (**see Table 2**). Motivated by the good metabolic stability we also evaluated the *in-vivo* pharmacokinetics of compound **10** in two different models (IV & PO) to evaluate its potential *in-vivo* application. Although the oral bioavailability for compound **10** was rather low (0.04) the MRTiv/po showed comparable results indicating its good metabolic stability (**see Table 2**).

**Table 2:** Pharmacokinetic evaluation of compound 10. Upper table: microsomal hepatic metabolic stability in human/mouse/rat; middle table: Mouse PK IV; lower table mouse PK PO.

| Microsomal Hepatic Metabolic Stability in |  |  |
| --- | --- | --- |
| Human | Mouse | Rat |
| 0.2mg/mL Clearance (uL/minute/mg) |  |  |
| 22.0 | 69.0 | 5.00 |

| Mouse PK IV |  |  |  |  |  |
| --- | --- | --- | --- | --- | --- |
| Cl<sub>total</sub><br>(mL/h/kg) | AUC<sub>iv</sub><br>(ng*h/mL) | C<sub>5min</sub><br>(ng/mL) | MRT<sub>iv</sub><br>(hours) | VD<sub>ss</sub><br>(mL/kg) | dose<br>(mg/kg) |
| 1056 | 94.7 | 44.5 | 1.59 | 1679 | 0.10 |

| Mouse PK PO |  |  |  |  |
| --- | --- | --- | --- | --- |
| AUC<sub>po</sub><br>(ng*h/mL) | C<sub>max</sub><br>(ng/mL) | MRT<sub>po</sub><br>(hours) | T<sub>max</sub><br>(hours) | dose<br>(mg/kg) |
| 30.9 | 11.8 | 1.78 | 0.50 | 1.0 |

### Conclusions

In this work, we present the discovery of the benzoxazepinone derivative **10** as a selective LIMK inhibitor based on a scaffold hopping approach. We have proven our estimation, namely that the specific αC-out / DFG-out seen in RIPK bound to benzoxazepinone derivatives can also be adapted by the highly flexible LIMK kinase domain and therefore be used to develop highly selective inhibitors targeting this unique conformation. The discovered compound exhibited low nanomolar potency, prolonged on-target residence time in KINETICfinder assays, strong cellular activity on the LIMKs in NanoBRET assays, and exceptional selectivity as shown by a comprehensive selectivity panel (KINOMEscan) without showing unspecific toxicity at recommended concentrations. Further, a phenotypic assay showed an effect of compound **10** towards cofilin phosphorylation and proved activity in a cellular environment. Considering target engagement characterization and off-target evaluation, we provide a novel LIMK1 and LIMK2 inhibitor with RIPK1 as single significant off-target and also a negative control **11** to profile our benzoxazepinone scaffold **10** as chemical tool compound. This compound will aid in clarifying the roles of LIMKs and their mechanisms of action in both pathogenesis and normal physiology.

## Methods

### Differential Scanning Fluorimetry Assay for LIMK1/2

The assay was performed according to a previously established protocol. A 2 μM solution of the respective LIMK in assay buffer (20 mM HEPES pH 7.4, 150 mM NaCl, 0.5 mM TCEP, 5% glycerol) was mixed 1:1000 with SYPRO Orange (Sigma-Aldrich). The compounds to be tested were added to a final concentration of 10 μM. Twenty microliters of each sample were placed in a 96-well plate and heated from 25 to 95 °C. Fluorescence was monitored using an Mx3005P real-time PCR instrument (Stratagene) with excitation and emission filters set to 465 and 590 nm, respectively. Data were analyzed with the MxPro software.

### DSF-based selectivity screening against a curated kinase library

The assay was performed as previously described.^29,35^ Briefly, recombinant protein kinase domains at a concentration of 2 μM were mixed with 10 μM compound in a buffer containing 20 mM HEPES, pH 7.5, and 500 mM NaCl. SYPRO Orange (5000×, Invitrogen) was added as a fluorescence probe (1 µl per mL). Subsequently, temperature-dependent protein unfolding profiles were measured using the QuantStudio™ 5 realtime PCR machine (Thermo Fisher). Excitation and emission filters were set to 465 nm and 590 nm, respectively. The temperature was raised with a step rate of 3°C per minute. Data points were analysed with the internal software (Thermal Shift SoftwareTM Version 1.4, Thermo Fisher) using the Boltzmann equation to determine the inflection point of the transition curve (**Table S4**).

### Isothermal Titration Calorimetry (ITC)

Measurements were performed at 20 °C on a MicroCal VP-ITC (GE Healthcare). LIMK1330⍰637 was dialyzed overnight into assay buffer (20 mM HEPES pH 7.4, 150 mM NaCl, 0.5 mM TCEP, 5% glycerol). The syringe was loaded with 105 μM LIMK1330⍰637, and the cell was filled with assay buffer containing 10 μM of the respective inhibitor. Every 5 minutes, 10 μL of the protein solution was injected into the cell for a total of 28 injections. The heat flow data were analyzed with the MicroCal ORIGIN software package employing a single binding site model.

### Protein Expression and Purification

The recombinant LIMK kinase domains LIMK1^330⍰637^ and LIMK2^330⍰632^ were expressed in insect cells and purified using affinity chromatography and size exclusion chromatography. In brief, exponentially growing TriEx cells (Novagen) at 2 × 10^6^ cells/mL were infected 1:64 with baculovirus stock, incubated for 66 h at 27 °C under constant shaking, and harvested by centrifugation. Cells were then resuspended in lysis buffer (50 mM HEPES pH 7.4, 500 mM NaCl, 20 mM imidazole, 0.5 mM TCEP, 5% glycerol) and lysed by sonication. The lysate was cleared by centrifugation and loaded onto a Ni^2+^ NTA column. After vigorous rinsing with lysis buffer, the His_6_-tagged proteins were eluted in lysis buffer containing 300 mM imidazole. While the proteins were subjected to dialysis to remove the imidazole, the N-terminal tags were cleaved by TEV protease. Contaminating proteins, the cleaved tags and TEV protease itself were removed with another Ni^2+^ NTA step. Finally, the LIMK kinase domains were concentrated and subjected to gel filtration using an AEKTA Xpress system combined with an S200 16/600 gel filtration column (GE Healthcare). The elution volumes of 91.8 mL (LIMK1^330⍰637^) and 91.6 mL (LIMK2^330⍰632^) indicated the proteins to be monomeric in solution. The final yields were 2.0 mg/L of insect cell medium for LIMK1^330⍰637^ and 0.2 mg/L for LIMK2^330⍰632^.

### Crystallization

One hundred nanoliters of drops of the protein solution with the respective ligand were transferred to a 3-well crystallization plate (SwisSci), mixed with 50 nL of the precipitant solution, and incubated at 4 °C (details in **Table S2**). Crystals appeared overnight and did not change appearance after 7 days. They were mounted with additional 25% ethylene glycol for cryoprotection. Data were collected at Diamond Light Source beamline I03, analyzed, scaled, and merged with Xia2. The structures were solved by molecular replacement with Phaser using a LIMK2 model as a template (PDB-ID 4TPT) and refined with Refmac5.^36^ The models were validated using MolProbity. The model and the structure factors have been deposited into the PDB (PDB ID: 8AAU).

### NanoBRET

The assay was performed as described previously.^29−31^ In brief, full-length LIMK1/2, RIPK1, BTK and LRRK2 cloned in frame in a NanoLuc-vector (Promega) were transfected into HEK293T cells (ATCC CRL-1573) using FuGENE HD (Promega, E2312), and proteins were allowed to express for 20 h. The serially diluted inhibitor and NanoBRET Kinase Tracer (Promega) used at the previously determined Tracer *K*_D_,app (**Table S1**) were pipetted into white 384-well plates (Greiner 781 207) using an ECHO 550 acoustic dispenser (Labcyte). The corresponding transfected cells were added and reseeded at a density of 2 × 105 cells/mL after trypsinization and resuspension in Opti-MEM without phenol red (Life Technologies). The system was allowed to equilibrate for 2 h at 37 °C and 5% CO2 prior to BRET measurements. To measure BRET, NanoBRET NanoGlo Substrate + Extracellular NanoLuc Inhibitor (Promega, N2160) were added as per the manufacturer′s protocol, and filtered luminescence was measured on a PHERAstar plate reader (BMG Labtech) equipped with a luminescence filter pair (450 nm BP, filter (donor) and 610 nm LP filter (acceptor)). Competitive displacement data were then plotted using GraphPad Prism 10 software using a normalized 3-parameter curve fit with the following equation: Y = 100/(1 + 10∧((X-Log IC50))).

### K192 NanoBRET selectivity screening

To assess the selectivity of compound **10**, the K192 Kinase Selectivity System (Promega, cat. no. NP4050) was used.^37^ For plate preparation, transfection mix was prepared in white 384-well small-volume plates (Greiner, cat. no. 784075) by pre plating 3 µL of 20 µL/mL FuGene HD (Promega, cat no. E2311), diluted in optiMEM medium (Gibco, cat. no. 11058-021). 1 µL DNA from both DNA vector source plates of the K192 kit was added using an Echo 550 acoustic dispenser (Beckman Coulter). The mix was incubated for 30 min and 6 µL HEK293T cells in optiMEM medium were added. The proteins were allowed to express for 20 h. After expression, Tracer K10 was added using the concentrations recommended in the K192 technical manual and 1 µM inhibitor was added to every second well. After 2 h of equilibration, detection was carried out using substrate solution comprising optiMEM with a 1:166 dilution of NanoBRET™ Nano-Glo® Substrate and a 1:500 dilution of the Extracellular NanoLuc® Inhibitor. 5 µL of substrate solution was added to every well and filtered luminescence was measured on a PHERAstar plate reader (BMG Labtech) equipped with a luminescence filter pair (450 nm BP filter (donor) and 610 nm LP filter (acceptor)). For every kinase, occupancy was calculated and plotted using GraphPad Prism 10 (**Table S5**).

### High-Throughput Kinetic Screening Assay

LIMK1 KINETICfinder^®^ assay was based on the binding and displacement of Alexa Fluor647-labeled kinase tracer to the ATP-binding site of the kinase with TR-FRET detection using terbium-labeled antibodies. The assays were performed in black 384 well microplates containing 0.1 nM of LIMK1 (Carna Biosciences), 30 nM of kinase tracer and 2 nM of Tb-labeled antibody (Life Technologies) in assay buffer (50 mM HEPES, pH 7.5, 10 mM MgCl_2_, 0.01% Brij-35, 1 mM DTT and 1% DMSO). For all experiments, a 4-point 10-fold serial dilution of 100X concentrated test compounds were prepared in DMSO. The kinetic assays were read continuously at room temperature in a PHERAstar FSX plate reader (BMG LABTECH). After collecting all the TR-FRET measurement, nonspecific TR–FRET signals were subtracted, and the specific signals were fitted to the Motulsky-Mahan’s “kinetics of competitive binding” equation. The affinity (K_d_), kinetic constants (k_on_, k_off_) and residence time of each test compound were calculated using KINPy® software (Enzymlogic).

### Multiplex High-content viability assessment

To assess the influence on cell health, a high-content screen in living cells called multiplex high content assay, as described previously by Tjaden et al. was performed.^38^ In brief, HEK293T (ATCC® CRL-1573™) and U2OS (ATCC®HTB-96™) were cultured in DMEM plus L-Glutamine (High glucose) supplemented by 10 % FBS (Gibco) and Penicillin/Streptomycin (Gibco). MRC-9 fibroblasts (ATCC® CCL-2™) were cultured in EMEM plus L-Glutamine supplemented by 10 % FBS (Gibco) and Penicillin/Streptomycin (Gibco). Cells were seeded at a density of 1250 cells per well in a 384 well plate in culture medium (Cell culture microplate, PS, f-bottom, µClear®, 781091, Greiner), with a volume of 50 µL per well. All outer wells were filled with 100 µL PBS-buffer (Gibco). Simultaneously with seeding, cells were stained with 60 nM Hoechst33342, 75 nM mitotracker red, 0.3 µL/well Annexin V Alexa Fluor 680 conjugate and 25 nL/well BioTracker™ 488 Green microtuble Cytoskeleton dye. Cell shape and fluorescence was measured before treatment, 12 h, 24 h and 48 h after compound treatment using the CQ1 high-content confocal microscope (Yokogawa, Musashino, Japan). The following setup parameters were used for image acquisition: Ex 405 nm/Em 447/60 nm, 500ms, 50%; Ex 561 nm/Em 617/73 nm, 100 ms, 40%; Ex 488/Em 525/50 nm, 50 ms, 40%; Ex 640 nm/Em 685/40, 50 ms, 20%; bright field, 300 ms, 100% transmission, one centered field per well, 7 z stacks per well with 55 µm spacing. The compounds were tested at 1 µM and 10 µM respectively. Acquired images of the cells were processed using the Yokogawa CellPathfinder software (v3.04.02.02). Cells were detected and gated with a machine learning algorithm as described previously (Tjaden et al. STAR protocols https://doi.org/10.1016/j.xpro.2022.101791). Results were normalized to cells exposed to 0.1 % DMSO. All compounds were tested in biological duplicates and SEM (standard error of mean) was calculated of biological duplicates. As a control compound for apoptotic cell death staurosporine (10 µM) was used (**Table S3**).

### Pharmacokinetic Screening

Test compounds were administered intravenously (0.1 mg/mL/kg) or orally (1 mg/5 mL/kg) by cassette dosing to non-fasted mice. After administration, blood samples were collected and centrifuged to obtain the plasma fraction. The plasma samples were deproteinized followed by centrifugation. The compound concentrations in the supernatant were analyzed by LC/MS/MS.

### Microsome Stability

Human liver microsomes were purchased from Sekisui Xenotech, LLC (Kansas city, KS). An incubation mixture consisted of microsomes in phosphate buffer (pH 7.4) and 1 µmol/L test compound. The concentration of microsomal protein was 0.2 mg/mL. An NADPH-generating system (MgCl_2_, glucose-6-phosphate, beta-NADP+ and glucose-6-phosphate dehydrogenase) was added to the incubation mixture with a half volume of the reaction mixture to initiate the enzyme reaction. The reaction was terminated 15 and 30 minutes after the initiation of the reaction by mixing the reaction mixture with acetonitrile, followed by centrifugation. The supernatant was subjected to LC/MS/MS analysis. The metabolic velocity was calculated as the slope of the concentration-time plot. The in vitro intrinsic metabolic clearance was calculated by dividing initial metabolic velocity by the test compound concentration in the incubation mixture.

### Chemistry

The synthesis of LIJTF500025 (**10**) and LIJTF500120 (**11**) was performed in WuXi AppTec Co., Ltd. The procedure and analytical characterization can be obtained from the supporting information.

### Treatment of cell lines and immunoblotting

LN-229, a human glioblastoma cell line, was cultured until passage 17 in Dulbecco’s modified Eagle’s medium (DMEM) with GlutaMAX^TM^, 10% fetal bovine serum (FBS), and 1% penicillin/streptomycin. Cells were maintained in an incubator with 5% CO_2_ at 37°C. For the treatment, 3 × 10^5^ cells were plated and treated with the tested compounds at the indicated concentrations and DMSO as the control for 6 hours. The cells were then lysed using RIPA buffer (50 mM Tris HCl pH 7.4, 150 mM NaCl, 1% Triton X-100, 1% NaDOC, 0.1% SDS, 1 mM EDTA), freshly supplemented with protease and phosphatase inhibitor cocktails (Sigma-Aldrich, Darmstadt, Germany). After a 1-hour incubation on ice, the lysates were centrifuged for 30 min at 250 × g, and the protein-containing supernatant was collected. The protein concentrations were determined using the Protein Assay Dye Reagent Concentrate (Bradford, Bio-Rad, Hercules, CA, USA) and Pre-Diluted Protein Assay Standards: Bovine Serum Albumin (BSA) Set (Thermo Fisher Scientific, Darmstadt, Germany). Then, equal amounts of protein were mixed with Roti®-Load 1 dye (Carl Roth, Karlsruhe, Germany) and 4x Laemmli Buffer (Bio-Rad, Hercules, CA, USA), denatured at 95 °C, and run on NuPAGE^TM^ 4−12% bis-Tris gels (Thermo Fisher Scientific, Darmstadt, Germany). Proteins were blotted on methanol-activated PVDF Transfer Membranes (Thermo Fisher Scientific, Darmstadt, Germany) using the wet transfer method with transfer buffer (Thermo Fisher Scientific, Darmstadt, Germany) and 20% methanol. Subsequently, the membranes were blocked with 5% milk or 3% BSA in 0.1% TBS-T for 1 h at room temperature and incubated overnight at 4 °C with primary antibodies: GAPDH (catalog no. 2118, Cell Signaling Technology, MA, USA, 1:1000 dilution), LIMK1 (catalog no. 3842, Cell Signaling Technology, 1:1000 dilution), LIMK2 (catalog no. 3845, 1:500 dilution), p-cofilin (catalog no. 3313, Cell Signaling Technology, 1:1000 dilution), cofilin (catalog no. 5175, Cell Signaling Technology, 1:1000 dilution). Then, the membranes were washed with 0.1% TBS-T and incubated with secondary horseradish-peroxidase (HRP)-conjugated antibody for 2 h (catalog no. 7074, Cell Signaling Technology, 1:3000 dilution). The membranes were washed again with 0.1% TBS-T before they were developed using X-ray films (Fujifilm, Dusseldorf, Germany).

## Supporting Information

The Supporting Information contains:

- Synthetic procedures; Figure S1 (ITC data), Figure S2 (Kinetic profiling using KINETICfinder), Table S1 (NanoBRET data), Table S2 (X-Ray data refinement).
- Table S3 (Multiplex cell-based assay data); Table S4 (DSF-Data); Table S5 (K192 NanoBRET selectivity screening); Table S6 (Eurofins Selectivity screening).

## Author Information

### Notes

The authors declare the following competing financial interest(s): B.-T.B. is a co-founder and the CEO of the Contract Research Organization CELLinib GmbH (Frankfurt am Main, Germany). The other authors declare no conflict of interest.

## Acknowledgements

The authors thank Mahmood Ahmed for his support to the project. The authors are grateful for support by the Structural Genomics Consortium (SGC), a registered charity (no: 1097737) that receives funds from Bayer AG, Boehringer Ingelheim, Bristol Myers Squibb, Genentech, Genome Canada through Ontario Genomics Institute [OGI-196], EU/EFPIA/OICR/McGill/KTH/Diamond Innovative Medicines Initiative 2 Joint Undertaking [EUbOPEN grant 875510], Janssen, Pfizer and Takeda. B-T.B. also received support by the collaborative research center CRC 1399 “Mechanisms of drug sensitivity and resistance in small cell lung cancer” and M.P.S. by the CRC1430 “Cell state transitions”. T.H. and S.K. are grateful for support by the ENABLE project “Unraveling mechanisms driving cellular homeostasis, inflammation and infection to enable new approaches in translational medicine”. A.K. S.K., M.P.S. and S.M. are also grateful for support by the translational cancer program of the German Cancer Aid TACTIC as well as by the Frankfurt Cancer Institute (FCI) and the translational cancer network DKTK.

## Abbreviations

LIMK1: LIM Domain Kinase1
RIPK1: receptor-interacting serine/threonine-protein kinase 1
TKL: tyrosine kinase like;

