## Supplemental Figure S1, Figure S2, Table S1, Table S2 for "Repurposing of the RIPK1 selective benzo[1,4]oxazepin-4-one scaffold for the development of a type-III LIMK1/2 inhibitor"

### Synthetical procedures:

#### Compound 15:

The synthesis was performed in WuXi AppTec Co., Ltd. According to the following protocols.

**Synthesis of ethyl 1-benzyl-5-hydroxy-1H-pyrazole-3-carboxylate (15):**

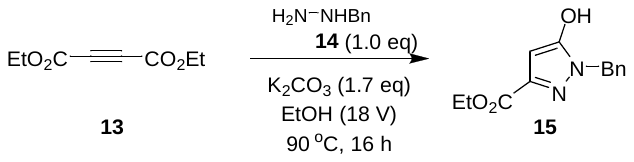

To a solution of K_2_CO_3_ (69.0 g, 499 mmol, 1.7 eq) and compound **14** (57.3 g, 293 mmol, 1.0 eq) in EtOH (830 mL) was added compound **13** (50.0 g, 294 mmol, 1.0 eq) at 15 °C. After addition, the mixture was stirred at 90 °C for 16 hours under N_2_ atmosphere. LCMS (product Rt = 0.844 mins) showed the reaction was completed. The reaction mixture was cooled to 15 °C, and was concentrated under reduced pressure to give a residue. The residue was diluted with H_2_O (250 mL) and extracted with MTBE (200 mL * 2). It was separated and the aqueous phase was adjusted with 6 M HCl to pH = 6, and it was extracted with ethyl acetate (200 mL * 3). The combined organic layer was washed with brine (200 mL), and concentrated under reduced pressure to give the crude product. The crude product was triturated with MTBE (20.0 mL) at 15 °C for 30 mins. And then filtered, the filter cake was washed with MTBE (20.0 mL). Compound **15** (25.0 g, 101 mmol, 99% purity, 34.2% yield) was obtained as a yellow solid.

^1^H NMR (400 MHz, CDCl_3_): δ 7.33-7.36 (m, 5H), 7.24-7.26 (m, 2H), 6.04 (s, 1H), 5.25 (s, 2H), 4.96 (s, 2H), 4.31-4.39 (m, 4H), 1.30-1.39 (m, 6H). LC-MS m/z [M + H]+: calcd 247.3, found 246.9.

#### Compound 16:

**Synthesis of ethyl 1-benzyl-5-bromo-4-formyl-1H-pyrazole-3-carboxylate (16):**

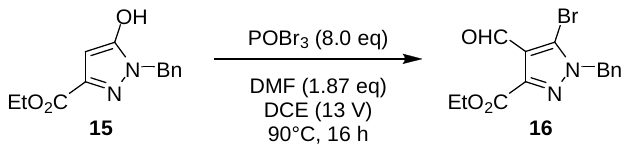

To a mixture of compound **15** (25.0 g, 102 mmol, 1.0 eq) and POBr_3_ (233 g, 812 mmol, 8.0 eq) in DCE (330 mL) was added DMF (13.9 g, 190 mmol, 14.6 mL, 1.87 eq) and the reaction was stirred at 90 °C for 16 hours under N_2_ atmosphere. TLC (petroleum ether/ethyl acetate = 5/1, product Rf = 0.30) indicated compound **15** was consumed completely. After cooling to room temperature, the mixture was added to ice water (800 mL) and it was extracted with DCM (800 mL * 2). The combined organic layer was washed with brine (600 mL) and dried over Na_2_SO4, then filtered and the filtrate was concentrated under reduced pressure. The residue was purified by column chromatography (SiO_2_, petroleum ether/ethyl acetate = 100/0 to 0/1). Compound **16** (7.40 g, 21.6% yield) was obtained as a yellow solid.

^1^H NMR (400 MHz, CDCl_3_): δ 10.4 (s, 1H), 7.26-7.46 (m, 5H), 5.51 (s, 2H), 5.25 (s, 2H), 4.46-4.51 (m, 2H), 1.44 (t, J = 7.2 Hz, 3H).

#### Compound 17:

**Synthesis of** **ethyl (*E*)-1-benzyl-5-bromo-4-(2-methoxyvinyl)-1H-pyrazole-3-carboxylate (17):**

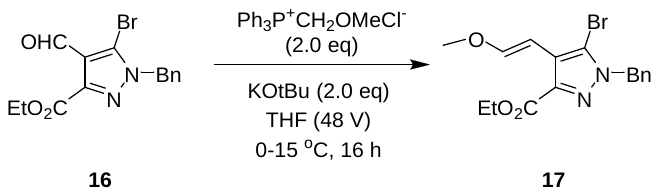

A mixture of Ph_3_P^+^CH_2_OMeCl^-^ (12.6 g, 36.7 mmol, 2.0 eq) in THF (150 mL) was added t-BuOK (4.12 g, 36.7 mmol, 2.0 eq) at 0 °C, and it was stirred at 0 °C for 0.1 hour. Then a solution of compound **16** (6.20 g, 18.4 mmol, 1.0 eq) in THF (150 mL) was added to the above solution at 0 °C, and the mixture was stirred at 15 °C for 16 hours under N_2_ atmosphere. LCMS (product Rt = 1.028 mins) showed compound **16** was consumed completely. The mixture was added to water (300 mL) and it was extracted with ethyl acetate (300 mL * 2). The combined organic layer was washed with brine (300 mL), dried over Na_2_SO_4_, filtered and the filtrate was concentrated under reduced pressure to give a residue. The residue was purified by column chromatography (SiO_2_, petroleum ether/ethyl acetate = 1/0 to 0/1). Compound **17** (4.00 g, 9.86 mmol, 53.6% yield, 90% purity) was obtained as a white solid.

^1^H NMR: (400 MHz, CDCl_3_) : δ 7.29-7.35 (m, 4H), 7.21-7.22 (m. 2H), 6.03 (d, *J* = 13.2 Hz, 1H), 5.45 (s, 2H), 4.41-4.46 (m, 2H), 3.71 (s, 3H), 1.43 (t, *J* = 7.2 Hz, 3H).

#### Compound 18:

**Synthesis of** **ethyl 1-benzyl-5-bromo-4-(2-oxoethyl)-1*H*-pyrazole-3-carboxylate (18):**

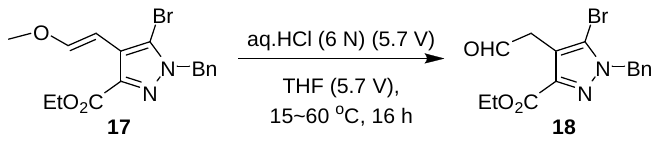

A mixture of compound **17** (3.70 g, 10.1 mmol, 1.0 eq) in THF (21 mL) was added HCl (6M, 21.0 mL, 12.5 eq) at 15 °C, and it was stirred at 60 °C for 1 hour. LCMS (product Rt = 0.973 mins) showed compound **17** was consumed completely. The reaction mixture was adjusted with sat.aq. Na_2_CO_3_ to pH = 7, and it was extracted with ethyl acetate (30.0 mL * 3). The combined organic layer was washed with brine (30.0 mL), dried over Na_2_SO_4_ and the filtrate was concentrated under reduced pressure to give a residue. The residue was purified by column chromatography (SiO_2_, petroleum ether/ethyl acetate = 1/0 to 0/1). Compound **18** (3.00 g, 7.60 mmol, 75.0% yield, 89% purity) was obtained as a colorless oil.

^1^H NMR (400 MHz CDCl_3_) : δ 9.61 (s, 1H), 7.17-7.36 (m. 5H), 5.42 (s, 2H), 4.31-4.36 (m, 2H), 3.74 (s, 2H), 1.32 (t, *J* = 7.2 Hz, 3H).

#### Compound 24:

**Synthesis of** **ethyl (*S*)-1-benzyl-5-bromo-4-(2-((5-methyl-4-oxo-2,3,4,5-tetrahydrobenzo[b][1,4] oxazepin-3-yl)amino)ethyl)-1*H*-pyrazole-3-carboxylate (20):**

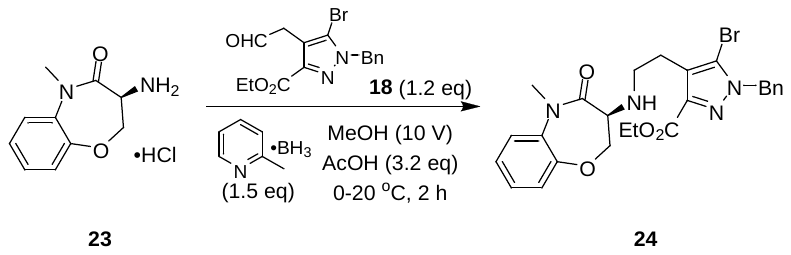

To a mixture of compound **18** (1.33 g, 3.78 mmol, 1.2 eq), AcOH (603 mg, 10.0 mmol, 3.19 eq) and compound **23** (0.72 g, 3.15 mmol, 1 eq, HCl) in MeOH (8.4 mL) was added borane-2-methylpyridine (491 mg, 4.59 mmol, 1.46 eq) at 0 °C and the mixture was stirred at 20 °C for 2 hours under N2 atmosphere. LCMS (product Rt = 0.874 mins) showed compound **23** was consumed completely. The reaction mixture was added to sat.aq. NaHCO_3_ (30.0 mL), and it was extracted with ethyl acetate (30.0 mL * 2). The combined organic layer was washed with brine (30.0 mL) and concentrated under reduced pressure. The residue was purified by prep-HPLC (column: Phenomenex Gemini-NX 80*40mm*3um; mobile phase: [water (10mM NH_4_HCO_3_)-ACN]; B%: 40%-70%, 8min). Compound **20** (700 mg, 1.33 mmol, 42.2% yield) was obtained as a yellow oil.

^1^H NMR (400 MHz CDCl_3_) : δ 7.31-7.32 (m, 6H), 7.16-7.27 (m, 2H), 5.42 (d, *J* = 15.2 Hz, 2H), 4.36-4.53 (m, 3H), 4.18 (s, 1H), 3.66-3.76 (m, 3H), 3.39 (s, 3H), 2.80-2.89 (m, 3H), 2.62 (s, 2H), 1.86 (s, 2H), 1.38 (t, *J* = 7.2 Hz, 3H).

#### Compound 25:

**Synthesis of** **(*S*)-3-(2-benzyl-3-bromo-7-oxo-2,4,5,7-tetrahydro-6*H*-pyrazolo[3,4-c]pyridin-6-yl)-5-methyl-2,3-dihydrobenzo[b][1,4]oxazepin-4(5*H*)-one (25):**

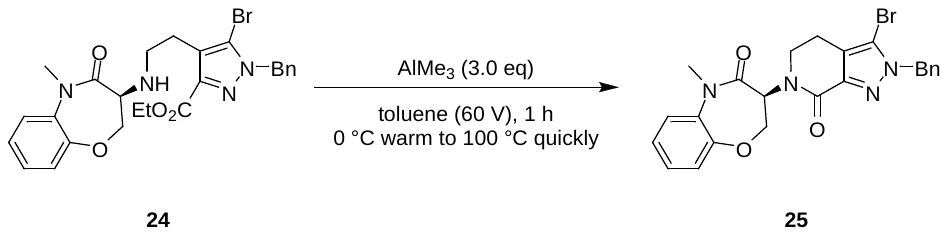

To a solution of compound **24** (600 mg, 1.14 mmol, 1.0 eq) in toluene (36.0 mL) was added Al(CH_3_)_3_ (2 M, 1.71 mL, 3.0 eq) at 0 °C, and the mixture was stirred at 100 °C for 1 hour. LCMS (product Rt = 0.874 mins) showed compound **24** was consumed completely. To the reaction mixture was added saturated aqueous potassium sodium tartrate solution (150 mL), and the mixture was stirred at room temperature for 30 mins, and extracted with ethyl acetate (60.0 mL * 2). The organic layer was washed with brine (50.0 mL), and dried over anhydrous Na_2_SO_4_, filtered and the filtrate was evaporated under reduced pressure. The residue was purified by column chromatography (SiO_2_, petroleum ether/ethyl acetate = 1000/1 to 1/1). Compound **25** (300 mg, 623 mmol, 54.8% yield) was obtained as a yellow solid.

^1^H NMR (400 MHz CDCl_3_) : δ 7.17-7.33 (m, 11H), 5.89-5.94 (m, 1H), 5.43 (s, 2H), 4.60-4.65 (m, 1H), 4.39-4.44 (m, 1H), 4.24-4.27 (m, 1H), 3.54-3.58 (m, 1H), 3.38 (s, 3H), 3.02-3.05 (m, 1H), 2.61-2.65 (m, 1H).

#### Compound 26:

**Synthesis of** **(*S*)-2-benzyl-6-(5-methyl-4-oxo-2,3,4,5-tetrahydrobenzo[b][1,4]oxazepin-3-yl)-7-oxo-4,5,6,7-tetrahydro-2*H*-pyrazolo[3,4-c]pyridine-3-carbonitrile (26)**

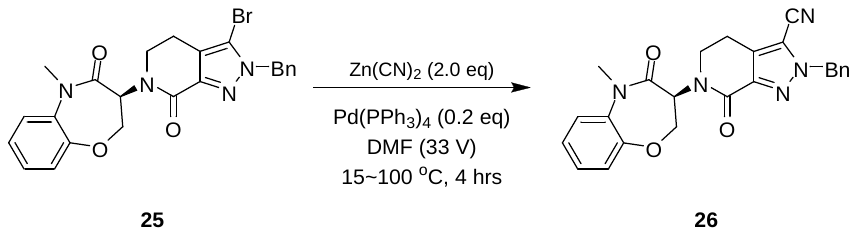

To a solution of compound **25** (300 mg, 623 umol, 1.0 eq) in DMF (10.0 mL) was added Pd(PPh_3_)_4_ (144 mg, 125 umol, 0.2 eq) and Zn(CN)_2_ (146 mg, 1.25 mmol, 2.0 eq) at 15 °C. The mixture was stirred at 100 °C under Ar for 4 hours. LCMS (product Rt = 0.874 mins) showed compound **25** was consumed completely. The mixture was quenched with water (50.0 mL) at room temperature and extracted with ethyl acetate 30.0 (mL * 2). The organic layer was separated, washed with brine 30.0 mL, dried over Na_2_SO_4_, filtered and the filtrate was concentrated in vacuo. The residue was purified by column chromatography (SiO_2_, petroleum ether/ethyl acetate = 1/0 to 0/1). Compound **26** (300 mg, 702 mmol, 84.4% yield) was obtained as a yellow solid.

#### Compound 27:

**Synthesis of** **(*S*)-2-benzyl-6-(5-methyl-4-oxo-2,3,4,5-tetrahydrobenzo[b][1,4]oxazepin-3-yl)-7-oxo-4,5,6,7-tetrahydro-2*H*-pyrazolo[3,4-c]pyridine-3-carboxamide (27):**

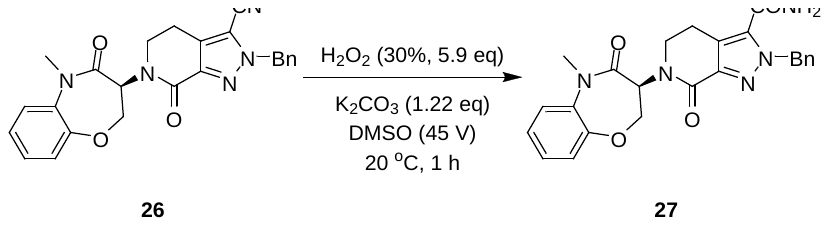

To a solution of compound **26** (220 mg, 515 umol, 1 eq) in DMSO (10.0 mL) was added K_2_CO_3_ (86.9 mg, 629 umol, 1.22 eq) and H_2_O_2_ (346 mg, 3.05 mmol, 5.93 eq, 30% w/w%) at 25 °C, and the mixture was stirred at 25 °C for 1 hour. LCMS (product Rt = 0.841 mins) showed compound **26** was consumed completely. The mixture was quenched with water at room temperature and extracted with ethyl acetate (30.0 mL * 3). The organic layer was separated, washed with sat. Na_2_SO_3_ (30.0 mL * 3) and brine, dried over Na_2_SO_4_, filtered and the filtrate was concentrated in vacuo. The residue was purified by column chromatography (SiO_2_, petroleum ether/ethyl acetate = 1/0 to 0/1). Compound **27** (180 mg, 384 umol, 74.6% yield, 95% purity) was obtained as a white solid.

^1^H NMR (400 MHz DMSO-d_6_) : δ 7.79 (s, 1H), 7.70 (s, 1H), 7.46-7.50 (m, 1H), 7.24-7.32 (m, 6H), 7.15-7.17 (m, 2H), 5.62 (s, 2H), 5.52-5.57 (m, 1H), 4.81-4.87 (m, 1H), 4.31-4.35 (m, 1H), 3.98-3.99 (m, 1H), 3.58-3.61 (m, 1H), 3.29 (s, 3H), 3.16 (d, *J* = 5.2 Hz, 1H), 3.03-3.04 (m, 1H), 2.80-2.86 (m, 1H).

#### Compound 28:

**Synthesis of (*S*)-6-(5-methyl-4-oxo-2,3,4,5-tetrahydrobenzo[b][1,4]oxazepin-3-yl)-7-oxo-4,5,6,7-tetrahydro-2*H*-pyrazolo[3,4-c]pyridine-3-carboxamide (28):**

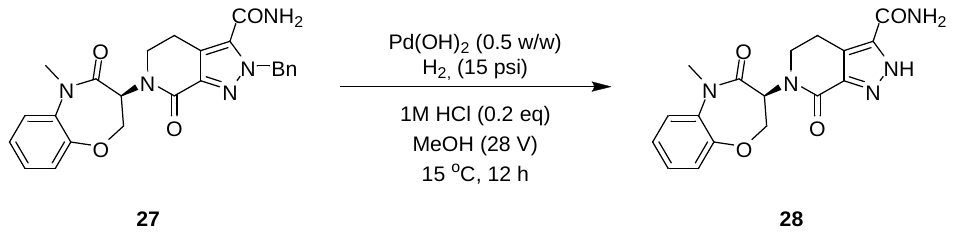

A mixture of compound **27** (180 mg, 404 umol, 1.0 eq), HCl (1 M, 80.8 uL, 0.2 eq), Pd(OH)_2_ (90.0 mg, 128 umol, 20% purity) in MeOH (5.00 mL) was degassed and purged with H_2_ for 3 times, and then the mixture was stirred at 15 °C for 12 hours under H_2_ at 15 psi. LCMS (product Rt = 0.516 mins) showed compound **27** was consumed completely. The reaction mixture was filtered through a pad of celite and the filtrate was concentrated in vacuo to give the residue. Compound **28** (150 mg, crude) was obtained as a brown solid and used for next step without purification.

#### Compound 11:

**Synthesis of (*S*)-1-benzyl-6-(5-methyl-4-oxo-2,3,4,5-tetrahydrobenzo[b][1,4]oxazepin-3-yl)-7-oxo-4,5,6,7-tetrahydro-1*H*-pyrazolo[3,4-c]pyridine-3-carboxamide (11):**

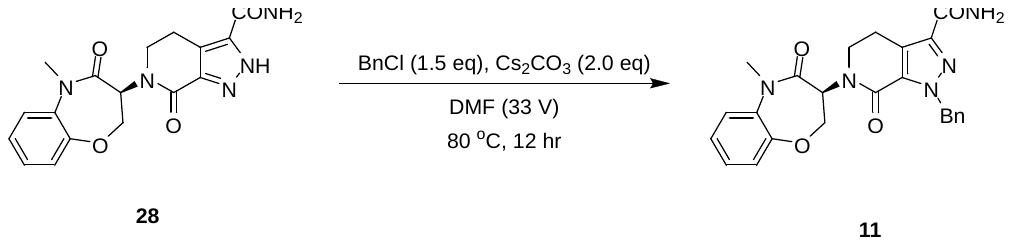

To a mixture of compound **28** (150 mg, 422 umol, 1.0 eq) in DMF (5.00 mL) were added Cs_2_CO_3_ (275 mg, 844 umol, 2.0 eq) and chloromethylbenzene (107 mg, 844 umol, 97.2 uL, 2.0 eq) slowly, and it was stirred at 80 °C for 4 hours. LCMS (product RT = 0.665 mins) showed compound **26** was consumed completely. The reaction mixture was concentrated under vacuum to give a residue. The residue was purified by prep-HPLC (column: Waters Xbridge BEH C18 100*30mm*10um; mobile phase: [water (10mM NH_4_HCO_3_)-ACN]; B%: 30%-50%, 10 min). Compound **11** (42.0 mg, 92.4 umol, 21.9% yield, 98.0% purity) was obtained as a white solid.

^1^H NMR (400 MHz DMSO-d_6_): δ 7.58 (s, 1H), 7.48 (d, *J* = 7.6 Hz, 1H), 7.25-7.31 (m, 7H), 7.17-7.18 (m, 2H), 5.62 (s, 2H), 5.46-5.51 (m, 1H), 4.83-4.88 (m, 1H), 4.33-4.38 (m, 1H), 4.01-4.02 (m, 1H), 3.64-3.66 (m, 1H), 3.30 (s, 3H), 3.01-3.05 (m, 2H).

#### Compound 20:

**Synthesis of ethyl (*S*)-1-benzyl-5-bromo-4-(2-((8-chloro-5-methyl-4-oxo-2,3,4,5-tetrahydrobenzo[*b*][1,4]oxazepin-3-yl)amino)ethyl)-1*H*-pyrazole-3-carboxylate (20):**

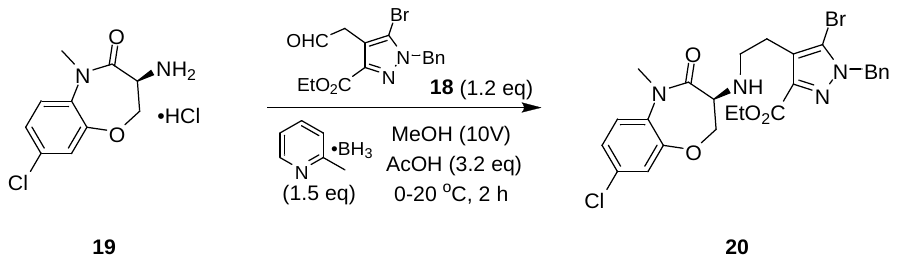

To a mixture of compound **18** (1.19 g, 3.37 mmol, 1.2 eq), AcOH (538 mg, 8.96 mmol, 3.19 eq) and compound **19** (740 mg, 2.81 mmol, 1 eq, HCl) in MeOH (7.40 mL) was added borane; 2-methylpyridine (438 mg, 4.10 mmol, 1.46 eq) at 0 °C and the solution was stirred at 20 °C for 2 hours under N_2_ atmosphere. LCMS (product Rt = 0.901 mins) showed compound **19** was consumed completely. The solution was added to sat. NaHCO_3_ (30.0 mL), and it was extracted with ethyl acetate (20.0 mL * 2). The combined organic layer was washed with brine (20.0 mL), filtered and the filtrate was concentrated under reduced pressure. The residue was purified by prep-HPLC (column: Phenomenex Gemini-NX 80*40mm*3um; mobile phase: [water (10mM NH_4_HCO_3_)-ACN]; B%: 30%-60%, 8 min). Compound **20** (800 mg, 1.42 mmol, 50.6% yield) was obtained as a yellow oil.

#### Compound 21:

**Synthesis of (*S*)-3-(2-benzyl-3-bromo-7-oxo-2,4,5,7-tetrahydro-6*H*-pyrazolo[3,4-*c*]pyridin-6-yl)-8-chloro-5-methyl-2,3-dihydrobenzo[*b*][1,4]oxazepin-4(5*H*)-one (21):**

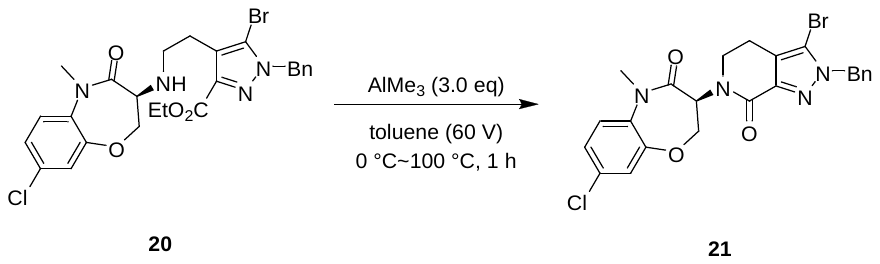

To a solution of compound **20** (570 mg, 1.01 mmol, 1.0 eq) in toluene (34.2 mL) was added Al(CH_3_)_3_ (2 M, 1.52 mL, 3.0 eq) at 0 °C, and the mixture was stirred at 100 °C for 1 hour. LCMS (product Rt = 1.024 mins) showed compound **20** was consumed completely. To the reaction mixture was added saturated aqueous potassium sodium tartrate solution (150 mL), and the mixture was stirred at room temperature for 30 mins, and extracted with ethyl acetate (80.0 mL * 2). The organic layer was washed with brine (60.0 mL), and dried over anhydrous Na_2_SO_4_, filtered and the filtrate was evaporated under reduced pressure. The residue was purified by column chromatography (SiO_2_, petroleum ether/ethyl acetate = 1000/1 to 1/1). Compound **20** (250 mg, 420 mmol, 41.4% yield) was obtained as a yellow solid.

^1^H NMR (400 MHz CDCl_3_) : δ 7.15-7.31 (m, 9H), 5.87-5.92 (m. 1H), 5.41 (d, *J* = 14.0 Hz, 2H), 4.60-4.66 (m, 1H), 4.39-4.43 (m, 1H), 4.21-4.25 (m, 1H), 3.52-3.54 (m, 1H), 3.35 (s, 3H), 3.00-3.05 (m, 1H), 2.61-2.64 (m, 1H).

#### Compound 22:

**Synthesis of (*S*)-2-benzyl-6-(8-chloro-5-methyl-4-oxo-2,3,4,5-tetrahydrobenzo[*b*][1,4]oxazepin-3-yl)-7-oxo-4,5,6,7-tetrahydro-2*H*-pyrazolo[3,4-*c*]pyridine-3-carbonitrile (22):**

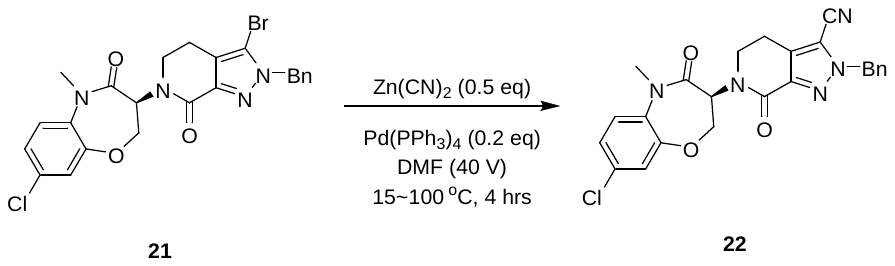

To a solution of compound **21** (250 mg, 485 umol, 1.0 eq) in DMF (10.0 mL) was added Pd(PPh_3_)_4_ (112 mg, 96.9 umol, 0.2 eq) and Zn(CN)_2_ (85.4 mg, 727 umol, 1.5 eq) at 15 °C. The mixture was stirred at 100 °C under Ar for 4 hours. LCMS (product Rt = 1.024 mins) showed compound **21** was consumed completely. The mixture was quenched with water (50.0 mL) at room temperature and extracted with ethyl acetate (50.0 mL * 2). The organic layer was separated, washed with brine (30.0 mL), dried over Na_2_SO_4_, filtered and the filtrate was concentrated in vacuo. The residue was purified by column chromatography (SiO_2_, petroleum ether/ethyl acetate = 1/0 to 0/1). Compound **22** (300 mg, 649 umol, 95.5% yield) was obtained as a yellow solid.

#### Compound 10:

**Synthesis of (*S*)-2-benzyl-6-(8-chloro-5-methyl-4-oxo-2,3,4,5-tetrahydrobenzo[*b*][1,4]oxazepin-3-yl)-7-oxo-4,5,6,7-tetrahydro-2*H*-pyrazolo[3,4-*c*]pyridine-3-carboxamide (10):**

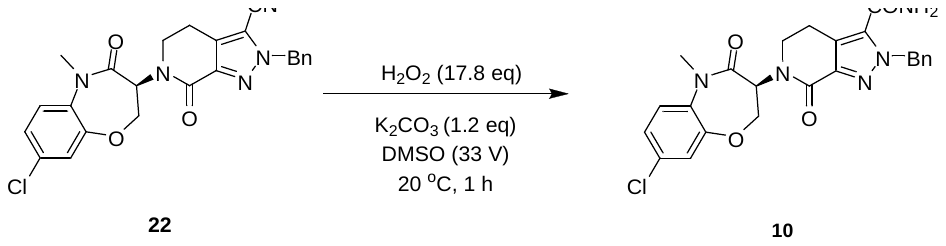

To a solution of compound **22** (300 mg, 649 umol, 1.0 *eq*) in DMSO (10.0 mL) was added K_2_CO_3_ (110 mg, 794 umol, 1.22 *eq*) and H_2_O_2_ (1.31 g, 11.56 mmol, 17.8 eq, 30% w/w%) at 25 °C, and the mixture was stirred at 25 °C for 1 hour. LCMS (product Rt = 0.925 mins) showed compound **22** was consumed completely. The mixture was quenched with water at room temperature and extracted with ethyl acetate (30.0 mL * 3). The organic layer was separated, washed with sat. Na_2_SO_3_ (30 mL * 3) and brine, dried over Na_2_SO_4_ and concentrated in vacuo. The residue was purified by prep-HPLC (column: Waters Xbridge BEHC18 100*30mm*10um; mobile phase: [water(10mM NH_4_HCO_3_)-ACN]; B%: 30%-60%, 10 min). **10** (31.0 mg, 64.6 umol, 9.95% yield, 100% purity) was obtained as a white solid.

^1^H NMR (400 MHz DMSO-d_6_): δ 7.82 (s, 1H), 7.72 (s, 1H), 7.55 (d, *J* = 8.8 Hz, 1H), 7.39 (d, *J* = 2.8 Hz, 2H), 7.30-7.32 (m , 3H),7.17-7.19 (m, 2H), 5.64 (s, 2H), 5.52-5.57 (m, 1H), 4.90 (t, *J* = 12 Hz, 1H), 4.38-4.43 (m, 1H), 3.97-4.00 (m, 1H), 3.60-3.62 (m, 1H), 3.29 (s, 3H), 3.03-3.05 (m, 1H), 2.86-2.88 (m, 1H).

### Supplementary Figures and Tables:

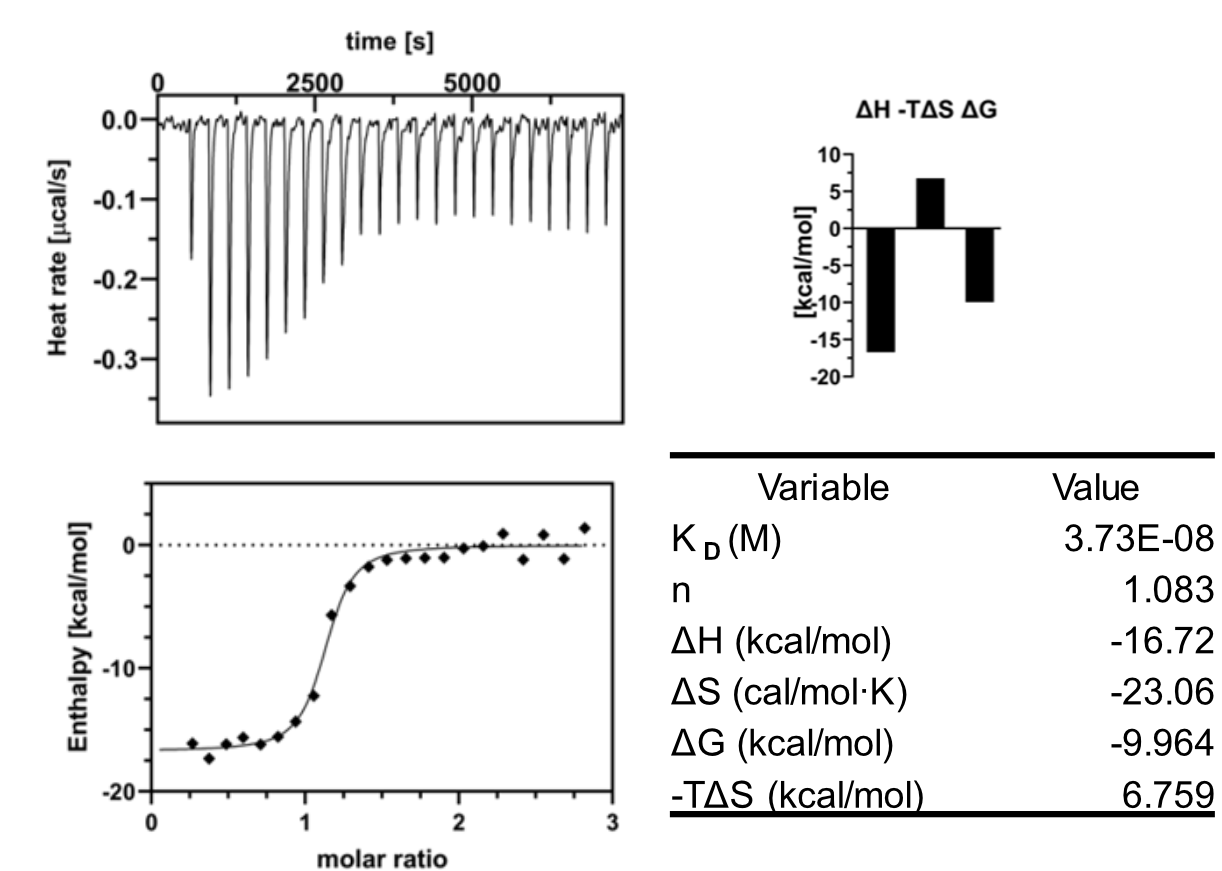

#### Figure S1: Isothermal Titration Calorimetry (ITC) of LIJTF500025a (10) on LIMK1.

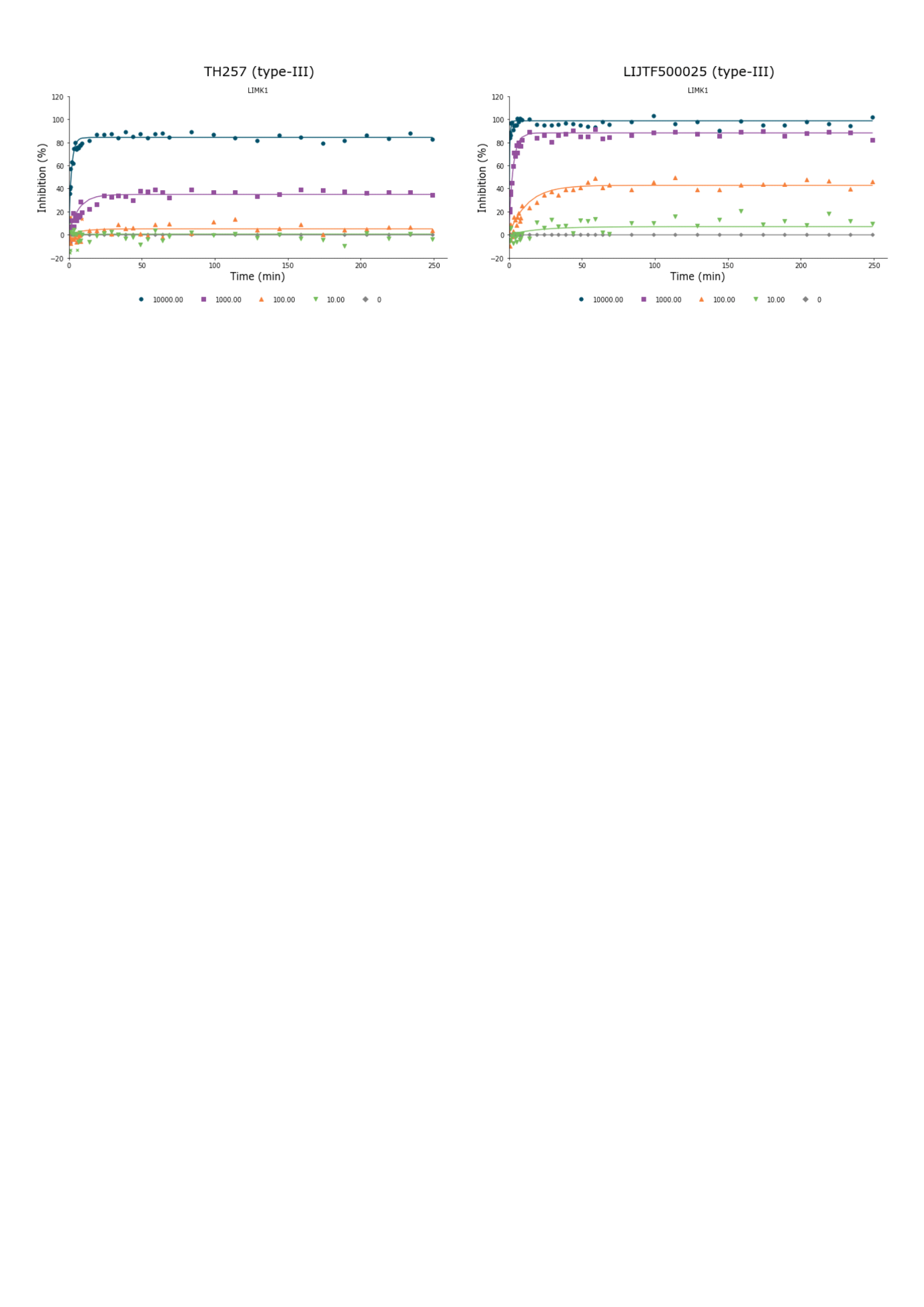

#### Figure S2: Kinetic profiling of TH257 (8) and LIJTF500025a (10) on LIMK1.

#### Table S1: NanoBRET data

| EC50 [M] | LIMK1 [nM] | LIMK2 [nM] | RIPK1 [nM] |
| --- | --- | --- | --- |
| LIJTF500025a | 82±6.5 | 52±6.3 | 6.3±0.23 |
| LIJTF500120a | > 50,000 | > 50,000 | 3,500±360 |
| TP-030-1 | 42,000±4,100 | 38,000±2,400 | 16±3.4 |
| TP-030-2 | 16,000±2,800 | 8,600±980 | 3.1±0.062 |

| **Protein** | **Plasmid catalog no. (Promega)** | **NanoLuc orientation** | **Tracer** | **Tracer catalog no. (Promega)** | **Tracer concentration used [M]** |
| --- | --- | --- | --- | --- | --- |
| LIMK1 | NV3391 | C | K10 | N2840 | 3.00E-07 |
| LIMK2 | NV1531 | C | K10 | N2840 | 4.00E-07 |
| RIPK1 | NV4171 | N | K10 | N2840 | 3.00E-07 |

#### Table S2. X-Ray Crystallography Data Collection and Refinement Statistics

|  | **LIMK1:LIJTF500025** |
| --- | --- |
| **PDB ID** | 7ATU |
| **Space group** | P 1 2_1_ 1 |
| **Cell parameters**  a, b, c (Å)  α, β, γ (°) | 85.23, 83.84, 98.03  90, 92.34, 90 |
| **Resolution (Å)** | 48.97 - 2.80 (2.94 - 2.80)* |
| **Unique reflexions** | 33787 (3396)* |
| **Completeness for range (%)** | 98.9 (99.7)* |
| **Multiplicity** | 3.3 (3.4) |
| **R_merge_** | 0.031 (0.348)* |
| **I/σ(I)** | 8.6 (1.8)* |
| **Wavelength (Å)** | 0.999 |
| **Phasing** | MR |
| **R_work_, R_free_ (%)** | 24.0, 31.6 |
| **Number of atoms**  protein, other, solvent | 7734, 136, 5 |
| **B-factors (Å^2^)**  protein, other, solvent | 62.7, 56.0, 41.6 |
| **rmsd bond (Å)** | 0.011 |
| **rmsd angle (°)** | 1.30 |
| **Ramachandran statistics**  favoured, outliers (%) | 89.2, 1.2 |
